# Ca^2+^ influx through ER-plasma membrane contacts is required for brown fat thermogenesis and metabolic health

**DOI:** 10.64898/2026.03.20.712802

**Authors:** J Zhou, C Dogan, LL Artico, K Rodrigues, S Hakam, T Kim, V Xu, K Lapenta, M Kang, DM Jorgens, SB Widenmaier, G Parlakgul, AP Arruda

## Abstract

Brown adipose tissue (BAT) exhibits exceptional metabolic plasticity, rapidly increasing energy expenditure to sustain thermogenesis during cold exposure. This high metabolic activity imposes substantial demands on cellular systems, requiring robust adaptive mechanisms to maintain homeostasis and prevent cellular stress. Yet, the pathways that support and coordinate these adaptive responses in brown adipocytes remain incompletely understood. Here, we identify a cold-induced adaptive program in BAT characterized by the formation of endoplasmic reticulum-plasma membrane (ER-PM) contact sites and the activation of store-operated calcium entry (SOCE), which is essential for maintaining brown adipocyte health during thermogenic activation. Cold exposure enhances ER-PM contacts and upregulates the expression of STIM and Orai proteins, key mediators of SOCE. Loss of STIM in brown adipocytes disrupts intracellular Ca²⁺ homeostasis and induces aberrant aggregation of ER membranes. STIM deficiency also impairs cold-induced mitochondrial fission resulting in hyperfused mitochondria with reduced oxidative capacity, independently of UCP1 abundance. Importantly, mice lacking STIM in BAT exhibit impaired lipid oxidation, are cold intolerant and develop exacerbated peripheral insulin resistance when challenged with a high-fat diet. Together, these findings identify ER-PM remodeling and STIM-mediated SOCE as a central regulator that links organelle architecture to brown adipocyte function and contributes to whole-body metabolic homeostasis.

## Introduction

Brown adipose tissue (BAT) exhibits a unique ability to promote futile nutrient oxidation and convert chemical energy into heat, making it a primary site of non-shivering thermogenesis^1,2^. The thermogenic capacity of brown adipocytes is largely attributed to their high mitochondrial content, uncoupling oxidative phosphorylation from ATP synthesis through UCP1 and the activation of futile cycles, including creatine and lipid cycling^3–7^. To maintain high oxidative activity, brown adipocytes rely on both intracellular and circulating lipids, glucose and branched-chain amino acids, thereby affecting systemic fuel levels^4,6,7^. BAT activity also impacts systemic metabolism by secreting a range of proteins, lipids, and metabolites that exert peripheral metabolic effects^8^. In a recent large-scale study, ^18^F-FDG PET scans revealed that individuals with detectable BAT had a lower prevalence of cardiometabolic diseases^9,10^. Conversely, reduced thermogenic capacity of adipocytes with age plays a key role in driving chronic diseases^11^. Thus, activation of BAT metabolic activity poses a potentially important therapeutic target for metabolic diseases.

Most research in the BAT field has focused on mechanisms that regulate mitochondrial metabolism and uncoupled respiration through UCP1^1,5,12^. However, organelles do not function in isolation. Emerging studies have revealed that other organelles, particularly the endoplasmic reticulum (ER), are also critical for maintaining the thermogenic capacity of BAT. For instance, cold exposure activates the ER-associated transcription factor NFE2L1, effecting proteasome activity and promoting protein quality control^13^. Disruption of this response leads to accumulation of ubiquitinated mitochondrial proteins, resulting in BAT dysfunction^13^. Similarly, recent studies have shown that inhibition of ER-associated protein degradation leads to cold intolerance^14^, and activation of PERK, a core component of the ER unfolded protein response, to regulate mitochondrial protein import and cristae remodeling in response to cold^15^. In addition, a futile Ca^2+^ cycling in the ER mediated by SERCA has been shown to induce thermogenesis independently of UCP1^16,17^. Thus, ER-related proteins and processes are critical to sustaining mitochondria homeostasis and metabolic activity of brown adipocytes.

In most cell types the ER is highly abundant, forming extensive physical contacts with all other organelles and the plasma membrane (PM) to coordinate the exchange of lipids, ions and metabolites^18,19^. Remodeling of ER inter-organelle physical communication is a central aspect of the ER’s adaptation in response to environmental changes, including nutritional challenges^19–21^. However, in brown adipocytes, the ER appears relatively sparse compared to the abundant mitochondria that dominate the cellular landscape^22^. How the ER is spatially organized in these cells and whether dynamic remodeling of ER structure and contact sites is required to support BAT functional adaptation to shifting metabolic demands remains poorly understood. This gap in knowledge is partly due to the technical challenges associated with high-resolution ultrastructural imaging of BAT, which has a uniquely complex subcellular architecture and is rich in lipid content.

Here, we show that cold exposure induces extensive remodeling of ER-PM contact sites in brown adipocytes, facilitating a previously unrecognized Ca^2+^ signaling mechanism in BAT via STIM-mediated store-operated Ca^2+^ entry (SOCE). While oscillations in cytosolic Ca²⁺ has been shown to modulate adrenergic signaling in brown adipocytes^23,24^, our findings identify SOCE at ER-PM contact sites as a distinct and essential mode of Ca²⁺ regulation required for remodeling of mitochondrial and ER structure and function during thermogenic activation. Importantly, loss of STIM-mediated SOCE in BAT uncouples adrenergic signaling from organelle dynamics, leading to impaired thermogenic capacity, cold intolerance, and insulin resistance during obesity.

## Results

### Cold exposure leads to ER-PM contact site remodeling as a previously unrecognized feature of BAT activation

To investigate how ER architecture adapts to thermogenic activation in vivo, we performed high-resolution 2D transmission electron microscopy (TEM) and 3D volume EM through Focused-Ion-Beam-Scanning Electron Microscopy (FIB-SEM) of murine brown adipose tissue. ER, mitochondria, lipid droplet, and nuclei were identified by manual labeling and automated segmentation using 3D-UNet-convolutional neuronal network models^25^. In mice housed at room temperature (RT), 2D TEM images depicted scarce ER either dispersed throughout the cytosol or associated with mitochondria and lipid droplets (**Fig. 1A)**. However, 3D reconstruction showed that the ER forms a network of membranes interspersed between the lipid droplets, mitochondria and other organelles (**Fig. 1B-C, video 1**). Distinct from other cells like hepatocytes^25^, brown adipocytes lacked parallelled stacks of rough ER sheets (**Fig. 1B, C**). The relative abundance of ER was around ∼1.5% of the cellular volume, whereas mitochondria and lipid droplets occupied about 20% and 38%, respectively (**Fig. 1D**).

**Figure 1.**
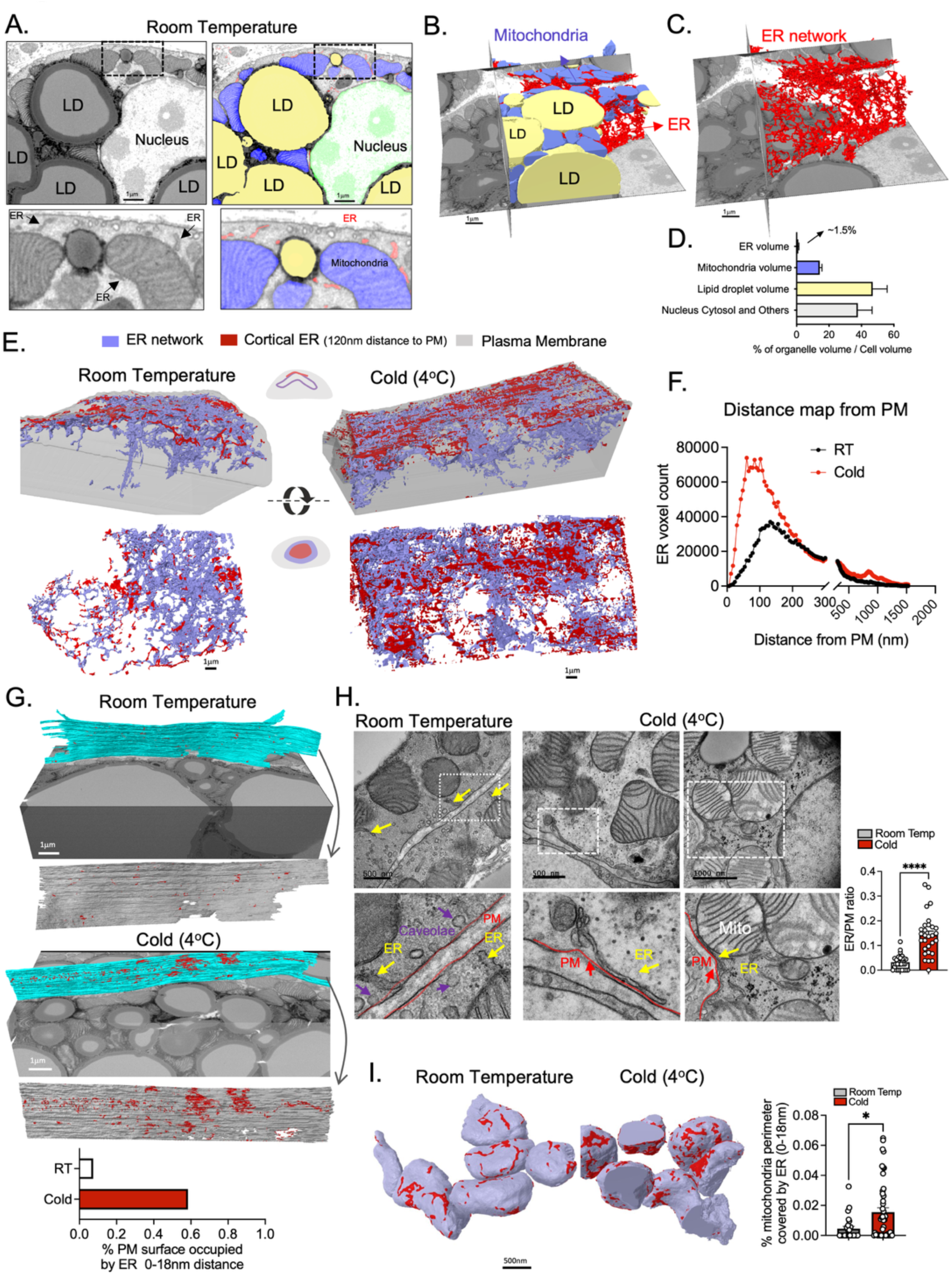
- Cold exposure induce ER-PM contact sites in brown adipocytes. (A) Left: Representative 2D scanning electron microscopy (SEM) image and Right: corresponding segmentation of brown adipose tissue (BAT) from C57BL/6J mice housed at room temperature (RT, 22 °C), highlighting subcellular structures. LD: lipid droplet; ER: endoplasmic reticulum; Mito: mitochondria. (B, C) 3D rendering of FIB-SEM volume showing LD (yellow), mitochondria (purple), and ER (red) of sample shown in A. (D) Quantification of organelle volumes normalized to cell volume in BAT from mice housed at RT. Data represent mean ± SEM from 3 different cells. (E) 3D rendering of the ER network reconstructed from BAT FIB-SEM images from mice housed at RT or exposed to cold (4°C) for 12 hrs. The ER network is shown in purple, with red highlighting the cortical ER located within 120 nm of the plasma membrane (PM). Upper panel: side view; Lower panel: front view; (F) Spatial distribution of ER voxel counts relative to PM distance, quantified from FIB-SEM datasets. (G) Upper panel: 3D rendering of PM (blue) highlighting areas of ER contacts within 18nm distance and Lower panel: quantification. (H) Left: Representative TEM images and Right: quantification (Right) from cortical areas of BAT tissue from mice maintained at RT or exposed to cold (4°C) for 12hrs. n= 28 images for RT and 29 images for cold, from 3 different mice per group. (I) Left: 3D rendering of FIB-SEM imaging and Right: quantification showing Mitochondria-ER contacts sites within 18nm distance (red). n= 25 mitochondria for RT; n=45 mitochondria for cold. The scale bar for all images is indicated in the figure. In all graphs data are represented as mean ± s.e.m. ****p< 0.001, * p<0.05, ** p<0.01 unpaired Student’s t-test. Reconstructions and analysis of FIB-SEM data were performed in Zeiss Arivis Pro software.

We then compared the subcellular organization of the ER and mitochondria in BAT isolated from mice housed at RT or exposed to cold (4°C) for 12 hours. Consistent with previous reports^26^, cold exposure induced mitochondrial fission, resulting in rounder organelles with increased circularity and reduced aspect ratio compared to the more elongated mitochondria present in BAT from mice at RT (**Extended Fig. 1A-D**). In cold-exposed BAT, the ER maintained its organization as a continuous network. However, in this condition, a significant fraction of ER membranes were redistributed toward cortical regions of the cell (**Fig. 1E, F, video 2,3**). Cold exposure also led to a marked increase in ER-PM contacts (∼18nm apart) in brown adipocytes (indicated in red in **Fig. 1G, video 4, 5**). Manual annotation of TEM images confirmed that ER-PM contact sites were rarely observed in mice housed at RT but became abundant in response to cold exposure (**Fig. 1H; Extended Fig. 1E-F**). In some cold-exposed cells, the ER formed tri-way junctions involving simultaneous contacts with the PM and mitochondria (**Fig. 1H; Extended Fig. 1F**). We further found that cold exposure not only reorganized ER-PM contacts, but also significantly increased ER-mitochondria interactions, as revealed by quantitative analysis (**Fig. 1I**), suggesting that in BAT, cold exposure significantly remodels organellar network interactions.

Cold-induced thermogenesis is driven by adrenergic signaling in response to norepinephrine (NE)^27,28^. To test whether adrenergic signaling in general drives ER remodeling toward the plasma membrane, we treated differentiated brown adipocytes with NE and measured ER-PM contact site abundance using MAPPER, a fluorescent reporter composed of an N-terminal ER-targeting signal, a transmembrane domain anchoring it to the ER, and a C-terminal polybasic motif that binds negatively charged PM phospholipids at a ∼25 nm spacing^29^ (**Extended Fig. 1G**). Confocal microscopy at the basal cell surface revealed that NE stimulation increased ER-PM contact sites, as indicated by enhanced MAPPER puncta formation (**Extended Fig. 1H**). Collectively, these findings demonstrate that cold exposure, likely through adrenergic stimulation, promotes ER remodeling, marked by increased ER-PM contact sites, alongside mitochondrial fission and enhanced ER-mitochondria interactions.

### Cold exposure leads to upregulation of STIM and Orai at ER-PM contact sites

ER-PM contact sites are key regulators of intracellular Ca^2+^ homeostasis through store-operated Ca^2+^ entry (SOCE)^18,30,31^. Given the prominent expansion of ER-PM contacts during cold exposure and the central role of Ca^2+^ signaling in brown adipocyte function^16,23,24,32^, we hypothesized that cold-induced ER remodeling allows SOCE activation to support Ca²⁺ homeostasis in conditions of increased thermogenic demand.

SOCE is mediated by the transmembrane ER protein and Ca^2+^ sensor STIM and the PM-localized Orai Ca^2+^ channels. Under resting conditions, STIM is distributed throughout the ER membrane and bound to Ca^2+^. Upon ER luminal Ca^2+^ depletion, STIM oligomerizes and translocates to ER-PM contact sites, where it interacts with Orai to promote Ca^2+^ influx from the extracellular space into the cytosol. This imported Ca^2+^ is then taken up into the ER to replenish Ca^2+^ stores and also coordinate Ca^2+^ signaling in the cytosol^18,30,31^. Two STIM isoforms (STIM1 and STIM2) and three Orai isoforms (Orai1-3) are expressed with tissue-specific patterns. Our analysis revealed that STIM1 is the main isoform expressed in adipose tissues and is enriched in BAT compared to other adipose depots (**Extended Fig 2A-B**). Orai1 and Orai3 are expressed in adipose tissues with Orai3 expression being higher in BAT compared to other depots, while Orai2 is not detected in any adipose depot (**Extended Fig 2C**). To our knowledge, this represents the first systematic characterization of STIM and Orai isoform expression across adipose depots, providing a molecular framework for understanding how SOCE is configured in thermogenic versus non-thermogenic adipocytes. Interestingly, both STIM and Orai mRNA and protein levels progressively increased in BAT as environmental temperature decreased (**Fig. 2A, B**). Cold exposure also induced STIM1 expression in inguinal white adipose tissue (iWAT) (**Extended Fig. 2D**), however in epididymal white adipose tissue (eWAT) its expression remains unchanged by temperature (**Extended Fig. 2E**).

**Figure 2.**
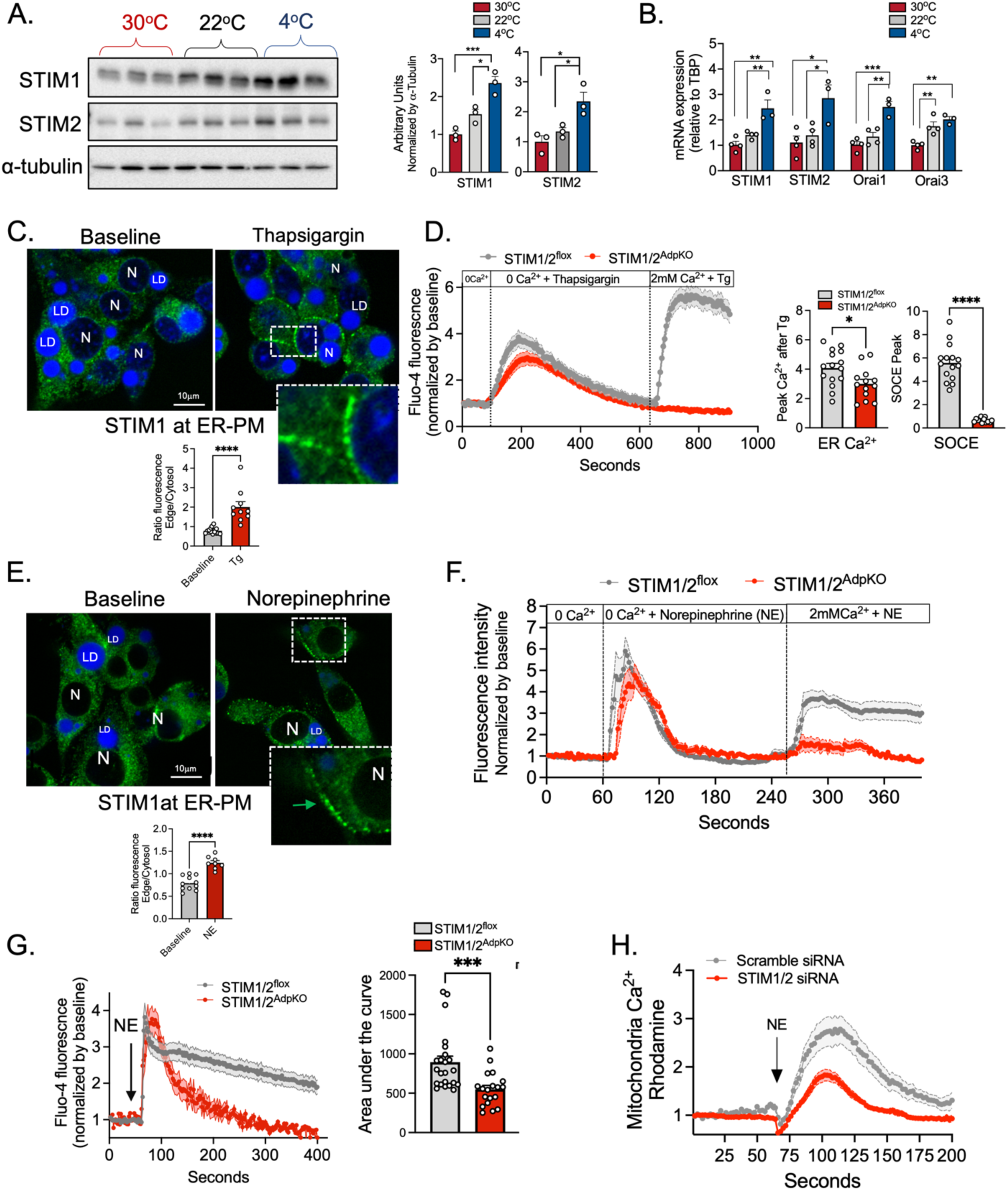
- Adrenergic stimulation leads to upregulation of STIM-Orai and activation of SOCE in brown adipocytes. (A) Left: Immunoblotting analysis of indicated proteins in total lysates from BAT of mice maintained at 30°C (thermoneutrality), 22°C (RT) or exposed to 4°C (cold) for 24hrs. Right: Quantification analysis of images shown in A; n= 3 mice per group. One-way ANOVA with Tukey’s multiple comparisons test, *p<0.05, *** p<0.001. (B) mRNA levels of indicated genes in BAT from mice exposed to 30°C, 22°C, n= 4 and 4°C for 24hrs, n=3. One-way ANOVA with Tukey’s multiple comparisons test, * p<0.05, ** p<0.01, *** p<0.005. (C) Upper panel: Immunostaining of STIM1 (green) in differentiated brown adipocytes in baseline and in cells treated with 1μM thapsigargin (Tg) for 10 min in Ca^2+^ free media; lipid droplets (LD) are stained in blue. Bottom: Quantification of the ratio of fluorescent intensity in the plasma membrane per intensity in the adjacent cytosolic area. n= 16 cells baseline, 10 cells Tg. Representative of 2 independent experiments; unpaired Student t-test ****p< <0.0001. (D) Left: Fluo-4 based cytosolic Ca^2+^ measurements in differentiated primary brown adipocytes from STIM1/2^flox^ (control) and STIM1/2^AdpKO^ mice. Cells were first treated with 1μM Tg in Ca^2+^ free media. Subsequently, 2mM Ca^2+^ was added in the indicated time point. The graph shows a representative run of 2 independent experiments. n=15 cells for STIM1/2^flox^ and n=13 cells for STIM1/2^AdpKO^. Right: Quantification of the cytosolic peak after Tg and Ca^2+^ addition. Unpaired Student t-test *p<0.05. (E) Upper panel: Immunostaining of STIM1 (green) in differentiated brown adipocytes in baseline and in cells treated with 10μM norepinephrine (NE) for 30 min in Ca^2+^ free media. Lipid droplets are stained in blue. Bottom: Quantification of the ratio of fluorescent intensity in the plasma membrane per intensity in the adjacent cytosolic area. n= 11 cells baseline, 8 cells NE. Representative of 3 independent experiments; Unpaired Student t-test ****p<0.0001. (F) Fluo-4-based cytosolic Ca^2+^ measurements in differentiated primary brown adipocytes from STIM1/2^flox^ (control) and STIM1/2^AdpKO^ mice. Cells were first treated with 10μM NE in Ca^2+^ free media. Subsequently, 2mM Ca^2+^ was added in the indicated time point. n= 10 cells per group; Representative of 3 independent experiments. (G) Left: Fluo-4-based cytosolic Ca^2+^ measurements in differentiated primary brown adipocytes from STIM1/2^flox^ and STIM1/2^AdpKO^ mice. Cells were treated with 10μM NE in the presence of 2mM Ca^2+^ in the media. The graph shows a representative run of 2 independent experiments. n=23 for STIM1/2^flox^, n=19 for STIM1/2^AdpKO^. Right: Area under the curve of the graph shown on the left. Unpaired Student t- test *** p<0.001. (H) Rhod-2-based mitochondria Ca^2+^ measurements in differentiated brown adipocytes transfected with scramble siRNA or with siRNA against STIM1/2. Cells were treated with 10μM NE in the presence of Ca^2+^ in the media. Representative of 2 independent experiments.

Beyond its role in SOCE, STIM1 also serves as an ER structural protein by interacting with the protein EB1 at the tip of the microtubules^33^. STIM1 overexpression has been shown to regulate cortical ER structure and increase the extent of ER-PM contact sites^34–36^. Therefore, cold-induced upregulation of STIM1 may contribute to the structural remodeling of ER-PM junctions. Consistent with this, expression of Junctophilin-2, a protein stabilizes ER-PM contacts in muscle^37^, was also higher in BAT of cold-exposed mice compared to controls (**Extended Fig. 2F**). These findings suggest that activation of BAT during cold not only enhances ER-PM contacts but coordinately upregulates the core machinery of the SOCE, positioning STIM-Orai signaling as a candidate mediator linking ER structural remodeling to Ca^2+^ homeostasis during BAT thermogenesis.

### Adrenergic signaling engages STIM-mediated SOCE shaping Ca^2+^ dynamics in brown adipocytes

Having established that cold exposure expands ER-PM and upregulates SOCE machinery, we next asked whether brown adipocytes are capable of engaging functional SOCE in response to adrenergic stimulation. We first assessed STIM1 localization in differentiated brown adipocytes treated with thapsigargin (Tg), a SERCA inhibitor that depletes ER Ca^2+^ stores and triggers STIM1 translocation to ER-PM contact sites. In both an immortalized brown adipocyte cell line as well as differentiated primary adipocytes (**Extended Fig. 2G, H**), Tg resulted in significant translocation of STIM1 to ER-PM junctions (**Fig. 2C and Extended Fig. 2I**). To directly measure SOCE, we performed single-cell Ca^2+^ imaging using a standard protocol involving ER Ca^2+^ depletion with Tg followed by re-addition of extracellular Ca^2+^ to monitor Ca^2+^ influx^34^. These experiments were conducted in differentiated primary adipocytes derived from both wild-type mice (STIM1/2^flox^) and mice with adipocyte-specific deletion of STIM1/2, using a Cre recombinase driven by the *Adipoq* promoter (STIM1/2^AdpKO^) (**Extended Fig. 2H**). Tg treatment, in the absence of extracellular Ca^2+^, elicited a sharp increase in cytosolic Ca^2+^ in both control and STIM1/2-deficient adipocytes, reflecting depletion of ER Ca^2+^ stores (**Fig. 2D**). However, the total amount of Ca^2+^ released from the ER was lower in STIM1/2-deficient cells compared to controls, suggesting that loss of STIM1/2 reduces steady-state ER Ca^2+^ levels (**Fig. 2D**). Subsequent addition of extracellular Ca^2+^ triggered Ca^2+^ entry in control cells, whereas this response was abolished in STIM1/2-deficient adipocytes, confirming that SOCE is STIM dependent in brown adipocytes (**Fig. 2D**).

Next, we tested whether adrenergic stimulation could regulate SOCE in physiologically relevant conditions. Treatment of brown adipocytes with NE in the absence of extracellular Ca^2+^ induced significant translocation of STIM to ER-PM contact sites (**Fig. 2E and Extended Fig. 2J**). NE stimulation also triggered an increase in cytosolic Ca^2+^, likely by activation ER Ca^2+^ release (**Fig. 2F**). Addition of Ca^2+^ in the media resulted in pronounced cytosolic Ca^2+^ increase in control cells, an effect that was reduced in adipocytes deficient for STIM1/2 (**Fig. 2F**). To more closely mimic physiological conditions, we acutely stimulated differentiated primary mouse brown adipocytes with NE in the presence of extracellular Ca^2+^. In both control and STIM1/2-deficient adipocytes, NE induced a rapid spike in cytosolic Ca^2+^ (**Fig. 2G**). However, whereas control adipocytes exhibited a sustained elevation of cytosolic Ca^2+^, STIM1/2-deficient cells showed a rapid return of cytosolic Ca^2+^ to baseline levels, as reflected by reduced area under the curve (**Fig. 2G**). Interestingly, we also found that downregulation of STIM1/2 resulted in reduced mitochondrial Ca²⁺ uptake in response to NE, as measured using the mitochondria-specific Ca²⁺ sensor Rhodamine-2 (**Fig. 2H**). These findings indicate that adrenergic stimulation activates SOCE in brown adipocytes, and that loss of SOCE alters ER, mitochondrial and cytosolic Ca^2+^ dynamics.

### Lack of STIM1/2 leads to defective cold-induced thermogenesis

To determine the physiological function of STIM-mediated SOCE in BAT, we examined the metabolic and thermogenic consequences of adipocyte-specific deletion of STIM1 and STIM2 in vivo (**Extended Fig. 3A**). STIM1/2^AdpKO^ mice showed decreased STIM1 and STIM2 protein expression in all adipose depots (**Fig. 3A; Extended Fig. 3B, C)**, with residual STIM expression likely due to contributions from non-adipocyte cell types present within the tissue.

**Figure 3.**
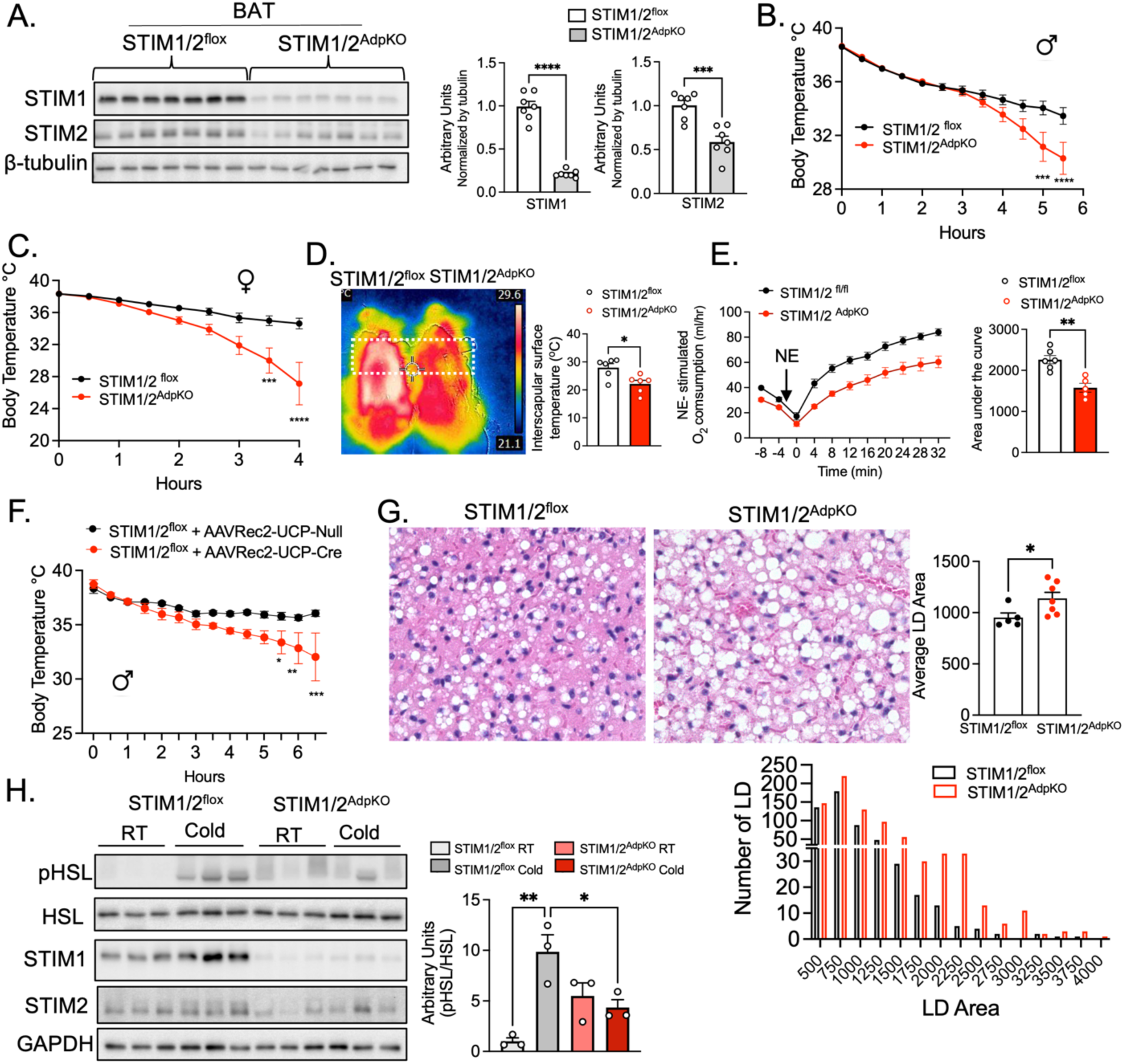
- Lack of STIM1/2 in adipocytes leads to impaired BAT lipid utilization and cold intolerance. (A) Left: Immunoblotting analysis of indicated proteins in total lysates of BAT from STIM1/2^flox^ (controls) and STIM1/2^AdpKO^. Right: Quantification of the images shown in the left. n=7 mice per group. Unpaired Student t-test, ****p<0.0001, *** p<0.005. (B, C) Body temperature over time of male (B) and female (C) mice exposed to 4°C in the absence of food. n =12 male mice per group and n =5 female mice per group. Two-way ANOVA with Šídák’s multiple comparisons test ***p<0.001,****p<0.0001. (D) Left: thermographic images from FLIR E5 infrared camera of STIM1/2^flox^ and STIM1/2^AdpKO^ mice exposed to 4°C for 4-6 hrs. Right: Quantification of images shown in the left. n= 6 mice per group. Unpaired Student t-test, *p<0.05. (E) Left: Norepinephrine-stimulated O2 consumption in anesthetized STIM1/2^flox^ and STIM1/2^AdpKO^ female mice. Right: Area under the curve of graph in the left. n= 6 mice STIM1/2^flox^ and n=5 mice STIM1/2^AdpKO^. Unpaired Student’s t-test ** p<0.001. (F) Body temperature over time of male STIM1/2^flox^ mice expressing AAV-Rec2-UCP1-Null AAV-Rec2-UCP1-Cre recombinase exposed to 4°C in the absence of food. n=4 Null mice and n=3 Cre mice per group. Two-way ANOVA with Šídák’s multiple comparisons test * p<0.05, ** p< 0.01, *** p=.0001. (G) Left: Representative Hematoxylin and eosin (H&E) stained sections of BAT from STIM1/2^flox^ and STIM1/2^AdpKO^ mice exposed to cold for 6hrs. Right: Quantification analysis of lipid droplet (LD) number and size of images shown in the left. n=5 STIM1/2^flox^, n=7 for STIM1/2^AdpKO^. Unpaired Student t-test * p<0.05. (H) Left: Immunoblotting analysis of indicated proteins in total lysates of BAT from STIM1/2^flox^ and STIM1/2^AdpKO^ maintained at RT or exposed to cold (4°C) for 2hrs. Right: Quantification of the images shown in the left. n=3 per group. One-way ANOVA with Tukey’s multiple comparisons test. * p<0.05, ** p< 0.005.

STIM1/2^AdpKO^ mice were born in normal Mendelian ratios and displayed similar body weights compared to their control counterparts (**Extended Fig. 3D**). However, when exposed to cold (4°C) both male and female STIM1/2^AdpKO^ mice became hypothermic after ∼ 4-5 hours of and had to be removed from cold chamber due to severe cold intolerance (**Fig. 3B, C**). Consistent with impaired BAT thermogenic activity, STIM1/2^AdpKO^ mice exhibited reduced BAT surface temperature during cold exposure, as measured by infrared thermo-imaging (**Fig. 3D**). To directly assess thermogenic capacity, we measured O2 consumption following subcutaneous NE injection above the interscapular BAT in anesthetized mice. NE-stimulated oxygen consumption was impaired in STIM1/2^AdpKO^ mice compared with controls (**Fig. 3E**). To determine whether this phenotype reflected a requirement for STIM specifically in brown adipocytes, we selectively downregulated STIM1/2 specifically in BAT using a recently described adeno-associated virus (AVV) serotype Rec2 which delivers Cre recombinase to BAT under the control of a mini-UCP1 promoter and simultaneously co-expresses a liver-targeted miRNA to suppress Cre expression in hepatocytes^38^ (**Extended Fig. 3E-F**). This approach efficiently deleted STIM1/2 in BAT without significantly altering STIM1/2 expression in other adipose tissue depots (**Extended Fig. 3G**). A cold tolerance test in this model showed that absence of STIM, specifically in BAT, leads to decreased cold tolerance similar to the Adipo-Cre model (**Fig. 3F**). To assess whether the cold intolerance observed in STIM1/2^AdpKO^ mice is linked to defective lipid mobilization, we performed histological analysis of BAT. Following cold exposure, STIM1/2^AdpKO^ mice displayed markedly larger lipid droplets compared to STIM1/2^flox^ control mice, suggesting impaired lipid utilization (**Fig. 3G**). A similar trend toward larger lipid droplets was observed at RT, although the difference did not reach statistical significance (**Extended Fig. 3H**). SOCE, activated downstream of STIM proteins, leads to a transient rise in cytosolic Ca^2+^ followed by ER Ca^2+^ refilling. Increased cytosolic Ca^2+^ is known to regulate lipolysis by modulating the activity of lipolytic enzymes^23,39^. Consistent with this, phosphorylation of hormone-sensitive lipase (HSL), an established marker of lipolytic activation, was reduced in STIM1/2^AdpKO^ mice upon cold exposure (**Fig. 3H**). These findings indicate that loss of STIM1/2 impairs Ca^2+^-dependent lipid mobilization in brown adipocytes.

### Lack of STIM leads to aberrant ER structure and function

To gain mechanistic insight into how SOCE regulates BAT function, we performed bulk RNA-seq transcriptomic analysis of BAT from STIM1/2^flox^ and STIM1/2^AdpKO^ mice at RT or exposed to cold. Under basal conditions, gene expression profiles were similar between the genotypes (**Fig. 4A**). In contrast, cold exposure for 6 hours revealed pronounced genotype-dependent transcriptional differences, with 589 genes differentially expressed in STIM1/2^AdpKO^ compared to controls, including 237 upregulated and 352 downregulated genes (**Fig. 4A, B**). Ingenuity pathway analysis of downregulated genes identified enrichment of pathways involved in oxidative stress response, protein ubiquitination, and hypoxia signaling pathways (**Fig. 4C**). Notably, the expression of Ppargc1a (PGC1α), a master regulator of BAT energy metabolism, was upregulated by cold exposure in control mice, but this induction was reduced in the absence of STIM1/2 (**Fig. 4B, D**). However, the expression levels of UCP1 were only slightly decreased in cold-exposed STIM1/2^AdpKO^ BAT (**Fig. 4B, D**), indicating that impaired thermogenesis is not primarily driven by loss of UCP1 expression. Among the top upregulated genes in BAT from cold-exposed mice lacking STIM1/2 were those involved in ER unfolded protein response (UPR) and quality control, such as HSPA5, Creld2 and Sdf2l1 (**Fig. 4B, E, F**). The UPR is mediated by three main branches - PERK, IRE1/XBP1, and ATF6 - which regulate overlapping transcriptional programs. Comparing our RNA-seq data with transcriptomic profiles from cells treated with selective activators of ATF6, IRE1, and PERK^40^ revealed that STIM deficiency preferentially activates an ATF6-driven transcriptional signature (**Fig. 4G**). Thus, loss of STIM triggers a distinct ER stress response in brown adipocytes likely due to altered ER Ca^2+^ homeostasis (**Fig. 2D**).

**Figure 4.**
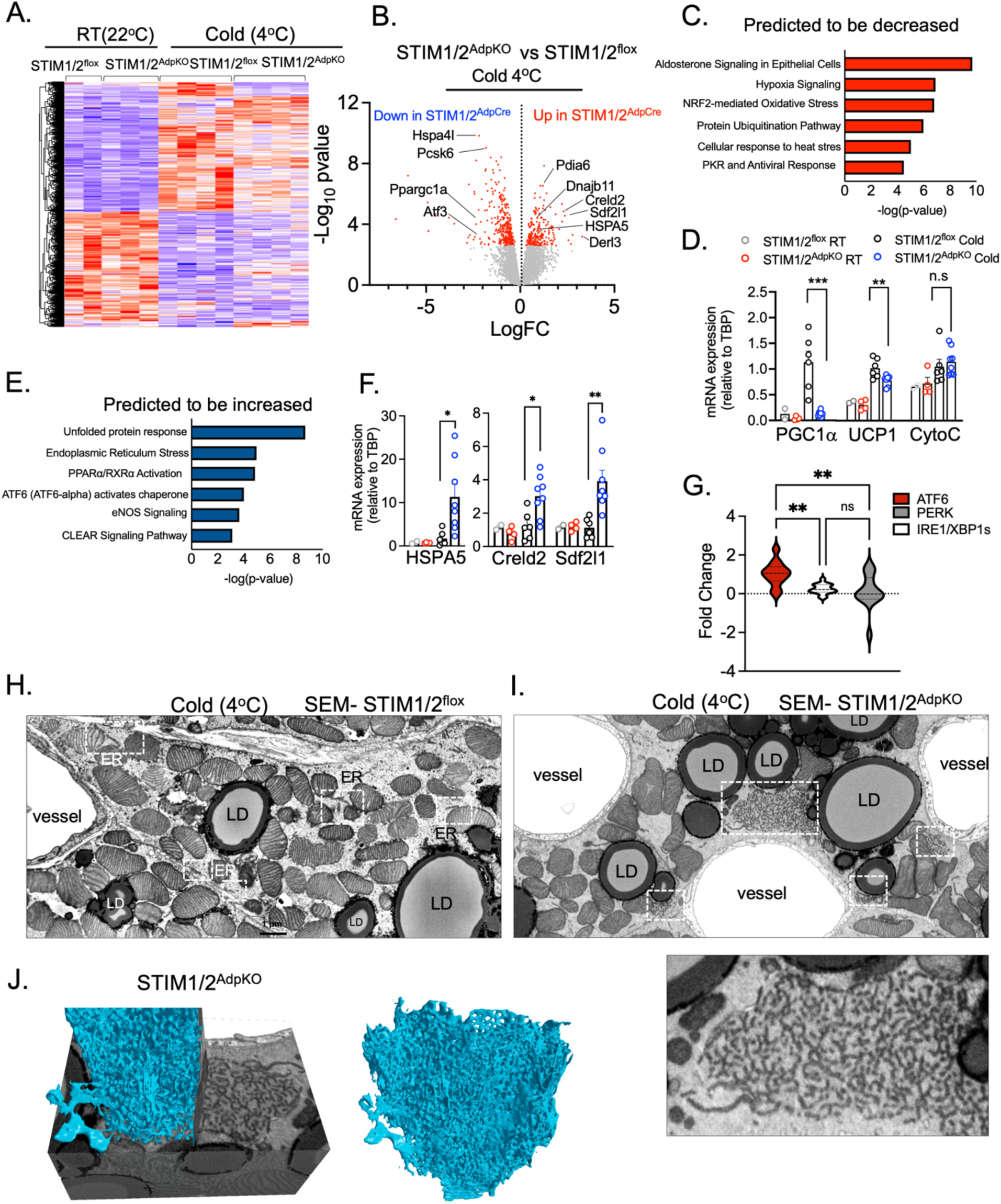
- Lack of STIM1/2 in adipocytes leads to increased ER stress and aberrant organelle morphology. (A) Heat map of RNA-seq analysis of mRNA expression of BAT from STIM1/2^flox^ and STIM1/2^AdpKO^ at RT or exposed to cold (4°C) for 6hrs; n= 2 mice for STIM1/2^flox^ RT; n=3 mice for STIM1/2^AdpKO^ RT; n= 4 mice for STIM1/2^flox^ and STIM1/2^AdpKO^ cold (B) Volcano plots showing differentially expressed genes comparing STIM1/2^flox^ and STIM1/2^AdpKO^ exposed to cold. Red dots show the significant regulated genes (p<0.05), Unpaired Student’s t-test. (C, E) Ingenuity pathway analysis showing pathways predicted to be decreased (C) or increased (E) in STIM1/2^AdpKO^ compared to STIM1/2^flox^ exposed to cold (4°C). p-value was calculated using Benjamin-Hochberg analysis. (D, F) qPCR-based analysis of mRNA expression of indicated genes in BAT from STIM1/2^flox^ and STIM1/2^AdpKO^ at RT or exposed to 4°C for 6h; n= 2 mice STIM1/2^flox^ RT; n = 4 mice STIM1/2^AdpKO^ RT; n = 5 mice STIM1/2^flox^ cold; n= 8 mice STIM1/2^AdpKO^ cold. Unpaired Student’s t-test *p<0.05, **p<0.01, ***p<0.0005. (G) Relative activation of ATF6, XBP1s and PERK gene sets from RNA seq analysis shown in A. These analyses were performed as previously described^40^. Differential activation of the ATF6, XBP1s, and PERK gene-sets was assessed by one-way ANOVA and significance of pairwise comparison confirmed by unpaired t-test. (H, I) SEM images of BAT tissue from STIM1/2^flox^ (H) and STIM1/2^AdpKO^ (I) mice exposed to cold (4°C) for 12hrs. White boxes highlight the ER/ aggregated membranes. (J) Segmentation and 3D reconstruction of FIB-SEM images of BAT from STIM1/2^flox^ and STIM1/2^AdpKO^ exposed to cold for 12hrs.

To determine whether the transcriptional changes were accompanied by alterations in ER structure, we performed TEM analysis of BAT tissue. At RT, the morphology of ER and other organelles were similar between the genotypes (**Extended Fig. 4A, B**), consistent with the minimal changes observed in the transcriptomic profile under these conditions (**Fig. 4A**). However, in response to cold, BAT tissues from STIM1/2^AdpKO^ mice showed a striking accumulation of aggregated membranes which were completely absent in the control (**Fig. 4H, I, video 6**). These aggregated membranes were located mostly in the cell periphery but were also detected in other cytosolic areas (**Fig. 4I; Extended Fig. 4C, D**). This phenotype was detected in multiple cells in BAT from different STIM1/2^AdpKO^ mice (**Extended Fig. 4C, D**). Volumetric FIB-SEM analysis showed that these aggregated membranes were continuous in the 3D, which suggests a tubular ER pattern (**Fig. 4J**). To our knowledge, these morphological alterations have never been detected in BAT before. Importantly, the aggregated membranes observed in the absence of STIM1/2 did not resemble the ER morphological changes seen in BAT under conditions of widespread ER stress induced by Tunicamycin in vivo, where the ER becomes markedly dilated (**Compare Fig. 4H-J and Extended Fig. 4E**). Thus, cold exposure in the setting of STIM1/2 deficiency activated a previously unrecognized ER adaptive response involving upregulation of ATF6-responsive genes and alterations in ER structural organization.

### STIM deficiency alters mitochondrial dynamics and decreases mitochondria oxidative capacity

During the analysis of the BAT 3D ultrastructure from cold-exposed control and STIM1/2^AdpKO^ mice, we noticed that in addition to the membrane aggregations described above, the mitochondrial morphology was also significantly altered in STIM1/2^AdpKO^ brown adipocytes. Ultrastructural analysis by FIB-SEM and TEM revealed that mitochondria from BAT of cold-exposed control mice appeared spherical (**Fig. 5A; Extended Fig. 5A, video 7**). However, mitochondria in STIM1/2^AdpKO^ adipocytes were significantly larger, less spherical, and exhibited a higher complexity index (**Fig. 5B-E, Extended Fig. 5A, video 8**). To further understand these findings, we examined mitochondrial dynamics in differentiated primary brown adipocytes from STIM1/2^flox^ and STIM1/2^AdpKO^ mice in response to adrenergic stimulation. Under basal conditions, mitochondria appeared elongated in both genotypes (**Fig. 5F, G**). Upon NE treatment, control primary brown adipocytes displayed a lower mitochondrial aspect ratio and reduced area, consistent with enhanced mitochondrial fragmentation (**Fig. 5F-H**). In contrast, in the absence of STIM1/2, mitochondria fail to undergo fission response upon NE stimulation. Some cells displayed mitochondrial morphology similar to untreated controls, while a subset (∼30%) of cells exhibited abnormally large, hyperfused mitochondria, indicative of impaired fission. (**Fig. 5F-H**). Thus, STIM1/2 deficiency in brown adipocytes impairs mitochondrial structural remodeling during thermogenic activation.

**Figure 5.**
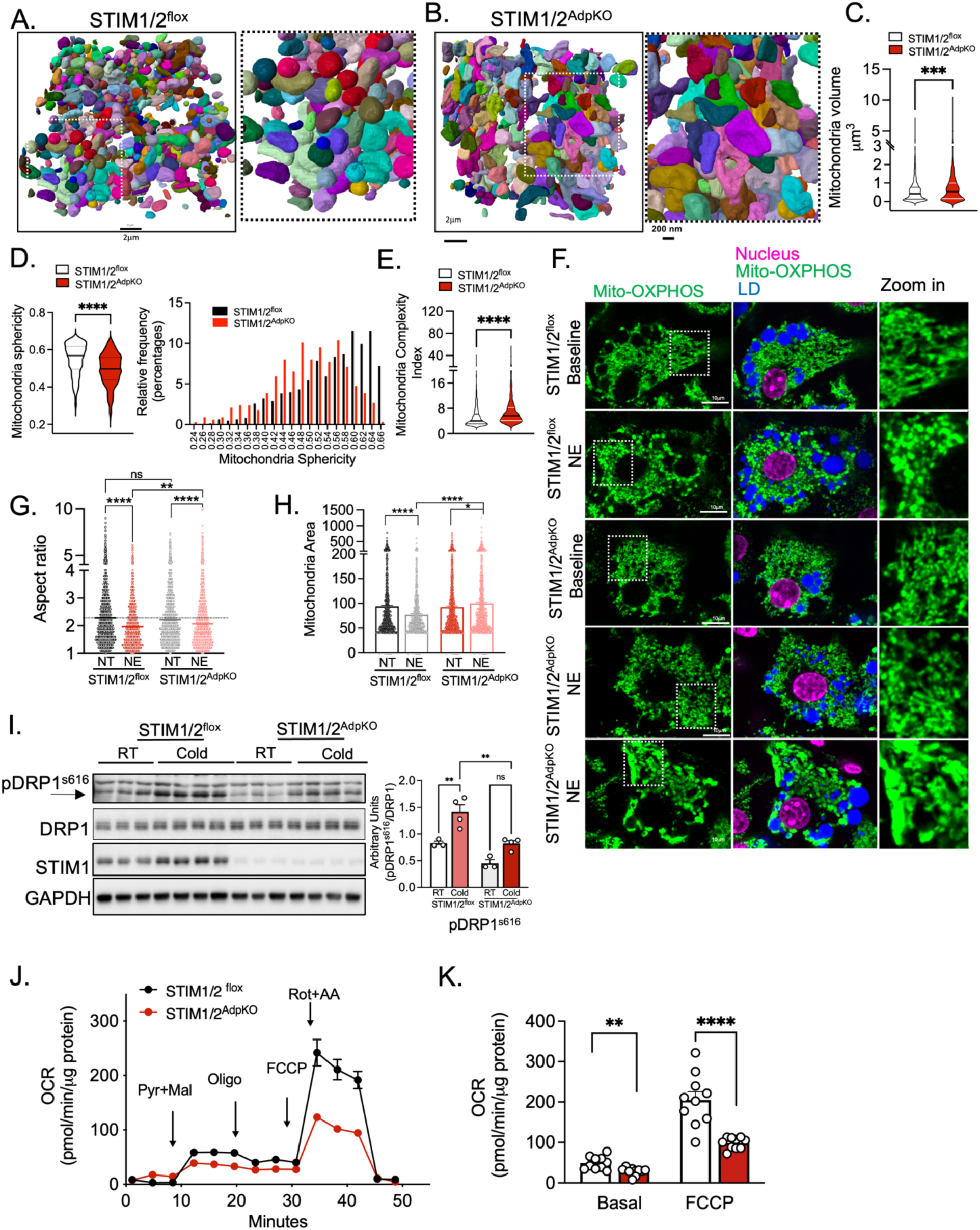
Lack of STIM1/2 in adipocytes lead to defects in ER and mitochondria structure. (A, B) Reconstruction of individual mitochondria from FIB-SEM data of BAT from STIM1/2^flox^ (A) and STIM1/2^AdpKO^ mice (B). 3D reconstructions were performed using Zeiss Arivis Pro software. (C) Quantification of mitochondria volume from data shown in A and B. n = 886 and 445 mitochondria, respectively. Unpaired Student’s t-test ***p<0.001 (D) Quantification of mitochondria sphericity and frequence distribution from data shown in A and B. n = 624 mitochondria in STIM1/2^flox^ and n=337 mitochondria in STIM1/2^AdpKO^. Unpaired Student’s t-test ****p<0.0001. (E) Mitochondria complexity index calculated according to^54^. Unpaired Student’s t-test ****p<0.0001. (F) Confocal images from differentiated primary adipocytes derived from STIM1/2^flox^ and STIM1/2^AdpKO^ mice at baseline or treated with 10uM norepinephrine (NE) for 50 min. Mitochondria (green) was staining with OXPHOs antibody, LD (blue) and Nucleus (magenta). (G,H) Quantification of mitochondria aspect ratio (G) and mitochondria area (H) of images shown in F. STIM1/2^flox^ NT= 1689 mitochondria, NE=861 mitochondria; STIM1/2^AdpKO^ NT=1185 mitochondria, NE=1618 mitochondria from > 5 images. Representative of 3 independent experiments. One-way ANOVA with Tukey’s multiple comparisons test **** p<0.0001. (I) Upper panel: Immunoblotting analysis of indicated proteins in total lysates of BAT from STIM1/2^flox^ and STIM1/2 ^AdpKO^ mice maintained at RT or exposed to cold (4°C) for 2hrs. Bottom panel: Quantification analysis of images shown in the upper panel. n=3 mice per group for RT conditions and n=4 mice per group for cold conditions. One-way ANOVA with Tukey’s multiple comparisons test. **p< 0.005. (J) Seahorse based oxygen consumption rate of mitochondria isolated from STIM1/2^flox^ and STIM1/2^AdpKO^ exposed to cold (4°C) for 12 hrs. Arrows depicts the addition of 20mM Pyruvate + 20mM Malate; 2μg/mL Oligomycin, 9μM FCCP and 4 µM rotenone and 2 µM antimycin. n=10 wells per group. Representative of 3 independent experiments. (K) Quantification of basal and maximal respiration. Unpaired Student t-test ** p<0.001, **** p<0.0001.

Mitochondrial dynamics (fission and fusion) are controlled by a coordinated network of dynamin-related GTPases^41^. Fusion of the outer mitochondrial membrane is mediated by Mitofusin 1 and Mitofusin 2, whereas mitochondrial fission is primarily driven by DRP1 in cooperation with its adaptor proteins MFF and FIS1. DRP1 is activated through post-translational modifications, most notably phosphorylation, which promotes its translocation to the outer mitochondrial membrane, where it assembles into oligomeric complexes to drive membrane scission^41^.

To determine whether the observed alterations in mitochondrial morphology stem from changes in the expression or activation of mitochondrial-shaping proteins, we examined their protein abundance and the phosphorylation status of DRP1. Mitofusin2 and MFF protein levels were not significantly changed in BAT lysates from STIM1/2^flox^ and STIM1/2^AdpKO^ mice exposed to cold (**Extended Fig. 5D)**. However, lack of STIM1/2 markedly affected the phosphorylation levels of DRP1. In BAT from STIM1/2^flox^ (controls), cold exposure resulted in increased phosphorylation of DRP1 in ser616; however, this induction was reduced in BAT from cold-exposed STIM1/2-deficient mice (**Fig. 5I)**. These findings suggest that STIM1/2 deficiency impairs mitochondrial fission, likely through reduced DRP1 activation. In addition, altered mitochondrial dynamics may also reflect the disrupted mitochondrial Ca^2+^ handling observed in **Fig. 2H**.

Given that mitochondrial morphology and Ca^2+^ handling are tightly coupled to bioenergetic function, we assessed mitochondrial function in BAT. We measured oxygen consumption in mitochondria isolated from BAT of STIM1/2^flox^ and STIM1/2^AdpKO^ mice exposed to cold using the Seahorse platform. Mitochondrial respiration was measured in the presence of guanosine diphosphate (GDP), an inhibitor of UCP1. Basal mitochondrial respiration in the presence of pyruvate and malate was slightly lower in STIM1/2^AdpKO^ BAT mitochondria compared with the controls (**Fig. 5I, J**). However, STIM1/2^AdpKO^ cells exhibited significantly higher respiratory capacity in response to maximal stimulation with FCCP, indicating impaired mitochondrial flexibility or reserve capacity (**Fig. 5I, J**). Importantly, these functional defects occurred without significant changes in the expression levels of electron transport chain subunits or UCP1 (**Extended Fig. 5B**), suggesting that the reduced respiratory performance is not due to altered abundance of key mitochondrial proteins. Altogether these findings show that STIM-mediated SOCE is required for proper mitochondrial structural remodeling and function during increased BAT metabolic demand.

### STIM is down regulated in obesity and STIM1/2 deficient mice show metabolic dysregulation

To determine the potential translational relevance of these findings, we examined the role of STIM1/2 in adipocytes in the context of obesity. Although BAT activation exerts broad metabolic benefits under physiological conditions^4,6,7^, chronic obesity is associated with reduced BAT mass and activity in both mice and humans^42,43^. To investigate whether altered STIM-mediated SOCE is involved in BAT dysfunction in obesity, we determined STIM1 and STIM2 expression in BAT from mice fed chow or a high-fat diet (HFD) for 16 weeks. As shown in **Fig. 6A**, STIM1 and STIM2 expression were significantly lower in BAT from HFD-fed obese mice compared to lean controls. To determine the impact of STIM deficiency on systemic metabolism in the context of obesity, we fed STIM1/2^flox^ and STIM1/2^AdpKO^ mice with HFD and examined how this intervention affected systemic metabolic parameters. STIM1/2^flox^ and STIM1/2^AdpKO^ gained body weight at a similar rate over the course of the HFD (**Fig. 6B**). However, when challenged with an insulin tolerance test, STIM1/2^AdpKO^ mice showed significantly higher insulin resistance compared to controls (**Fig. 6C**). Increased insulin resistance was also observed by oral glucose tolerance test, since to maintain similar glucose excursion rates, STIM1/2^AdpKO^ mice released significantly higher levels of insulin (**Fig. 6D, E**). Histological analysis revealed that BAT in HFD-fed STIM1/2^AdpKO^ mice accumulate much larger lipid droplets compared to BAT from STIM1/2^flox^ mice (**Fig. 6F**). To determine whether these effects were specifically due to STIM loss in BAT rather than other adipose depots, we performed similar experiments in mice in which STIM1/2 were selectively depleted in BAT via AAV-Rec2-UCP1-Cre delivery, following the protocol outlined in **Fig. 6G**. This approach effectively deleted STIM in BAT (**Fig. 6H**), while other adipose depots were unaffected (**Extended Fig. 6A, B**). Selective loss of STIM in BAT during HFD feeding led to impaired insulin sensitivity, as evidenced by reduced glucose clearance during both an insulin tolerance test (**Fig. 6I**) and glucose tolerance test (**Fig. 6J**). Together, these findings demonstrate that STIM1/2 is suppressed in BAT during obesity and that STIM-mediated Ca^2+^ signaling in brown adipocytes is required to preserve BAT metabolic function and systemic insulin sensitivity under conditions of diet-induced obesity.

**Figure 6.**
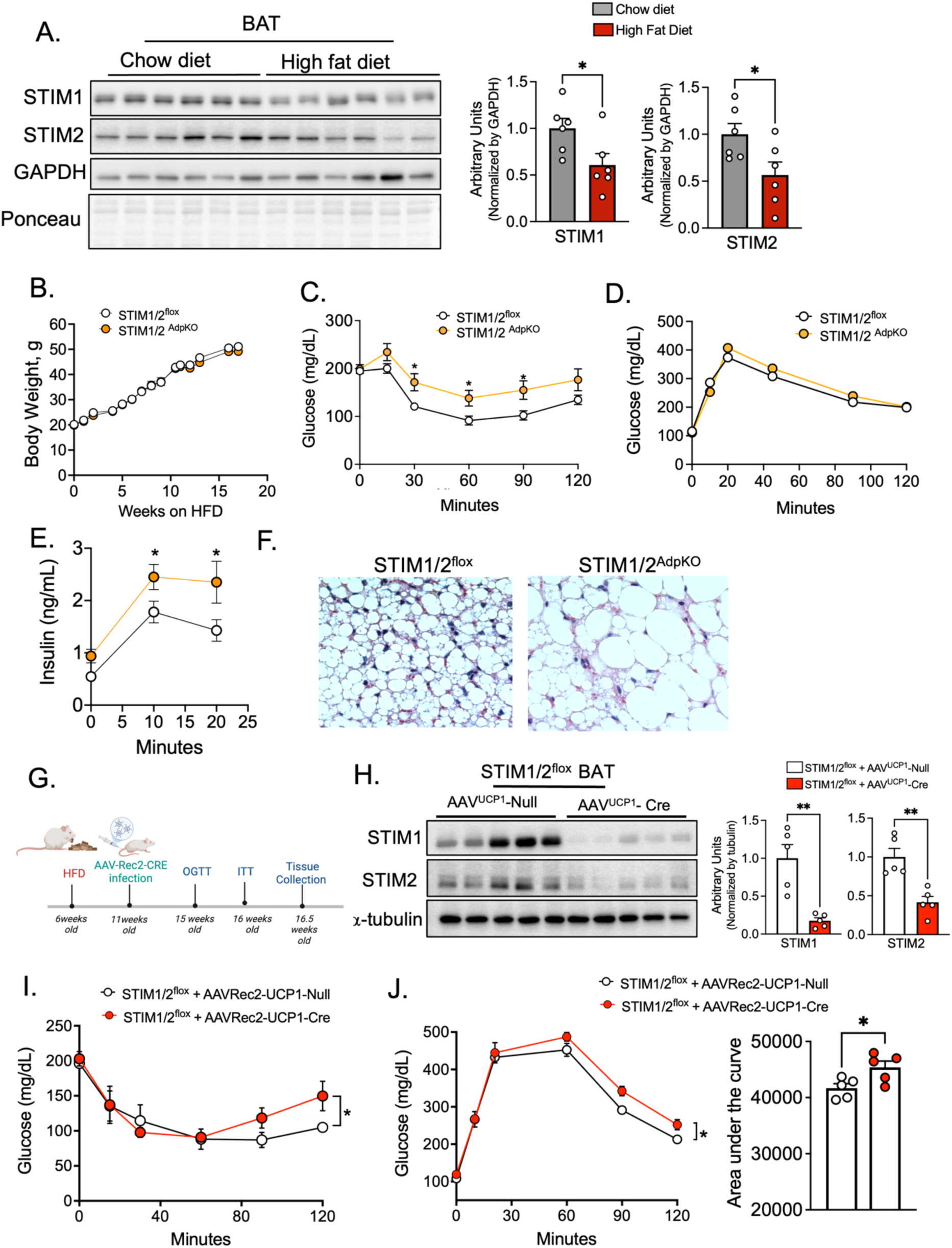
- Lack of STIM1/2 in adipocytes worsens insulin sensitivity in obese mice. (A) Left: Immunoblotting analysis of indicated proteins in total lysates from BAT of mice fed chow diet of high fat diet for 16 weeks. Right: Quantification analysis of the images shown in A; n= 6 mice for chow diet, n=6 mice for HFD group. Unpaired Student’s t-test, *p<0.05 (B) Body weight gain of STIM1/2^flox^ and STIM1/2^AdpKO^ mice fed a HFD over the indicated time. n=16 STIM1/2^flox^ mice, n=10 STIM1/2^AdpKO^ mice. (C) Insulin tolerance test, n =15 STIM1/2^flox^ mice, n=10 STIM1/2^AdpKO^ mice fed a HFD for 10 weeks. Two-way ANOVA **p < 0.01. (D) Oral glucose tolerance test, n=16 STIM1/2^flox^ mice, n=10 STIM1/2^AdpKO^ mice fed a HFD for 11 weeks (E) Insulin levels during the oral glucose tolerance test shown in D, Two-way ANOVA *p <0.05. (F) Representative hematoxylin and eosin-stained histology sections of BAT from STIM1/2^flox^ and STIM1/2^AdpKO^ mice fed HFD for 14 weeks. (G) Schematic depicting protocol of high fat diet (HFD) feeding and AAV-Rec2-UCP1-Null or - AAV-Rec2-UCP1-Cre injection. (H) Left: Immunoblotting analysis of indicated proteins in total lysates from BAT of mice fed HFD infected with AAV-Rec2-UCP1-Null or - AAV-Rec2-UCP1-Cre. Right: Quantification analysis of the images shown in A; n= 5 mice per group. Unpaired Student’s t-test, **p<0.01. (I) Insulin tolerance test, n= 4 Null mice and n=5 Cre mice; Two-way ANOVA *p <0.05. (J) Left: Oral glucose tolerance test, n =5 mice in each group. Two-way ANOVA *p <0.05. Right: Area under the curve from the graph shown on the left. Unpaired Student’s t-test *p<0.05

## Discussion

Remodeling of ER contact sites is increasingly recognized as a key regulator of cellular metabolism, facilitating adaptation to environmental changes through the regulation of intracellular Ca^2+^ transport, lipid exchange, organelle dynamics, and other mechanisms^18,19^. However, most studies have relied on immortalized cell lines, limiting our understanding of how these structures function in specialized tissues such as adipose tissue in a physiological context. Our previous work characterized the 3D structural organization of ER in liver tissue across fasting, fed, and obese states, revealing that ER morphology and its inter-organelle interactions are highly sensitive to metabolic conditions. This dynamic remodeling is disrupted in obesity, impairing tissue’s ability to adapt to metabolic demands^20,25^.

In this study, we leveraged high-resolution 3D volume EM and TEM to resolve organelle morphology in BAT, providing the first high resolution 3D ultrastructural characterization of ER morphology in this tissue. Our findings reveal that dynamic remodeling of ER-PM contact sites during cold exposure is essential for precise cytosolic and inter-organellar Ca^2+^ handling, which in turn sustains signaling pathways that regulate substrate utilization and organelle structure and dynamics (**Extended Fig.7**). These mechanisms are central for enabling brown adipocytes to meet the elevated metabolic demands imposed by cold exposure and contribute to insulin sensitivity in the context of obesity.

Intracellular Ca^2+^ signaling is tightly controlled in space and time, coordinating a wide range of cellular responses. In brown adipocytes, activation of Ca^2+^ influx through the plasma membrane L-type voltage-gated Ca^2+^ channels in response to adrenergic stimulation leads to rise in cytosolic Ca^2+^ and activate downstream PKA signaling^24,44,45^. In this study, we demonstrate that intracellular Ca^2+^ regulation in brown adipocyte response to adrenergic stimulation is more complex than previously anticipated. Adrenergic signaling induces ER Ca^2+^ release which subsequently activates SOCE, resulting in a sustained elevation of cytosolic Ca^2+^ levels, followed by ER Ca^2+^ replenishment. Disruption of this regulatory system markedly alters the kinetics of cytosolic Ca²⁺ signaling, reduces steady-state ER Ca²⁺ storage, and limits Ca²⁺ uptake by mitochondria. These changes impair lipid mobilization and compromise mitochondrial structure and function, at least in part by disrupting the proper activation of kinase signaling pathways, including pHSL and pDRP1 (**Extended Fig.7**). Additionally, reduced ER Ca^2+^ levels resulting from impaired SOCE trigger a distinct ER stress response which seems to be preferentially driven by ATF6, consistent with previous findings showing that ATF6 activation compensate for ER stress induced by reducing luminal Ca^2+^ level^46,47^.

A striking structural consequence of STIM1/2 loss in BAT was the marked accumulation of membrane aggregates upon cold challenge. Based on their 3D continuity with the ER, we hypothesize that these structures represent reorganized ER membranes. This reorganization does not appear to result from increased phospholipid biosynthesis, as genes involved in lipid synthesis pathways were not upregulated in this setting. Interestingly, similar membrane aggregates are observed in skeletal muscle affected by tubular aggregate myopathy, a disorder caused by mutations in STIM1^48,49^. Given that brown adipocytes and skeletal muscle share a common developmental origin^50^, the phenotype we observe in STIM1/2-deficient BAT may be mechanistically analogous to the tubular aggregates seen in STIM1-linked myopathy.

Another key finding of our study is that the loss of STIM-mediated SOCE disrupts mitochondrial morphology, Ca^2+^ handling and oxidative capacity in activated brown adipocytes. Given the lack of large changes in the expression levels of mitochondrial including UCP1 and ETC components, these observations suggest that the reduced mitochondrial oxidative capacity is not due to impaired biogenesis or expression of core metabolic genes but rather may result from a failure to undergo proper organelle structural remodeling in response to adrenergic stimulation. Altered mitochondrial Ca²⁺ homeostasis may further contribute to this phenotype. Indeed, increased Ca²⁺ uptake through the mitochondrial calcium uniporter (MCU) is known to stimulate TCA cycle activity, enhance NADH production, and promote mitochondrial respiration, indicating that impaired Ca²⁺ transfer to mitochondria could directly limit oxidative metabolism^32^.

Importantly our findings may also have implications for metabolic disease. In humans, obesity is associated with reduced BAT activity but the mechanisms behind this observation remain largely unknown. We found that obesity in mice downregulates STIM1/2 expression in BAT, and that deletion of STIM1/2 in brown adipocytes impairs systemic insulin sensitivity in the context of obesity, highlighting a previously unrecognized role for SOCE in BAT-mediated metabolic regulation. These results identify SOCE as an important component of BAT-mediated metabolic regulation and suggest that impaired Ca²⁺ signaling may contribute to BAT dysfunction during metabolic disease.

In conclusion, our study identifies a novel mechanism regulating brown adipocyte metabolism that integrates Ca^2+^ signaling with cold-induced dynamic organelle structural remodeling. We demonstrate that STIM-mediated Ca^2+^ signaling is essential for maintaining brown adipocyte integrity, enabling adaptive mitochondrial and ER remodeling, and supporting systemic metabolic homeostasis in response to both cold exposure and chronic high-fat feeding. Together, these findings position SOCE as a key regulator of brown adipocyte function and a potential target for improving metabolic health.

## Materials and Methods

### General animal care

All experimental procedures involving animals were approved by the University of California Berkeley Institutional Animal Care and Use Committee (Protocol# AUP-2022-02-15079-1 approved in May 2022). All mice were housed in groups (3-4 mice per cage) at 22 ± 1°C with bedding in the cage on a 12h light/dark cycle with free access to water and standard laboratory chow diet (PicoLab Mouse Diet 20 #5053) in the UC Berkeley pathogen-free barrier facility. For cold exposure (4°C) experiments, mice were single housed in cages with bedding with ad libitum access to drinking water and chow diet. In some experiments food was removed during cold exposure as stated in the legend of the figures.

#### Adipocyte-specific STIM1/2 loss of function mice

Mice carrying floxed alleles for STIM1 and STIM2 on a C57BL/6J background were kindly provided by Dr. Patrick Hogan from the La Jolla Institute for Allergy and Immunology. To generate adipocyte-specific STIM1/2 deficient mice, STIM1^flox^ mice were bred with STIM2^flox^ mice to generate double floxed mice and subsequently bred with mice expressing CRE recombinase under the control of the adiponectin promoter (AdpCre) in C57BL/6J background (Jackson Labs stock #028020). Age-matched littermates were used for the study. For high fat diet (HFD) studies, mice were fed HFD (D12492i: 60% kcal% fat; Research Diets) starting at 6 weeks of age for up to 15-18 weeks. The control group remained on chow diet (PicoLab Mouse Diet 20 #5053). Number of replicates and sample size were determined based on prior research^13,51^.

#### Brown adipocyte-specific STIM1/2 loss of function mice

To achieve brown adipocyte-specific deletion of STIM1/2, we used a recombinant adeno-associated virus (rAAV) serotype Rec2 expressing Cre recombinase under the control of a mini-UCP1 promoter and co-expressing microRNA-122. The virus was administered intravenously via tail vein injection in STIM1/2^flox^ mice at a dose of 3×10^12^ or 1×10^13^ IFU in 100 μL of saline per mouse. Control mice received an identical AAV construct lacking the Cre recombinase gene. The rAAV-Ucp1-Cre and -null control vectors were kindly provided by Dr. Ludger Scheja (University Medical Center Hamburg-Eppendorf, Germany; see^38^ for vector details). The pseudotyped Rec2 AAV vectors were a gift from Dr. Deborah Young (University of Auckland, New Zealand). Vector generation and titration were performed by Franklin BioLabs Inc. For the HFD studies, STIM1/2^flox^ mice were first fed a HFD for 5 weeks before receiving either rAAV-Ucp1-Cre or control AAV. Mice remained on the HFD for an additional 4-5 weeks, during which metabolic testing was conducted. Tissue collection occurred 5 weeks post-injection, as indicated. Mice were euthanized by cervical dislocation and organs designated for RNA, lipid, and protein analyses were snap-frozen in liquid nitrogen.

### Transmission electron microscopy (TEM) analysis

#### Sample preparation and TEM image acquisition

TEM sample preparation was performed as previously described^25^. Briefly, mice were perfused with saline through the heart, followed by a fixative containing 2.5% glutaraldehyde and 2.5% paraformaldehyde in 0.1 M sodium cacodylate buffer (pH 7.4) (Electron Microscopy Sciences, catalog #15949). After perfusion, small biopsies of brown adipose tissue were collected and further incubated in a fresh fixative containing 1.25% formaldehyde, 2.5% glutaraldehyde, 0.03% picric acid, and 0.05 M sodium cacodylate buffer. Samples were embedded in Epon resin and ultrathin sections were prepared using a Reichert Ultracut-S microtome. Samples were imaged with a Tecnai 12 electron microscope at 80 kV.

#### TEM image data analysis

For the data presented in **Fig. 1H** and Extended **Fig. 1C, D**, TEM images at the same magnification were manually annotated in FiJi to identify plasma membrane, ER, and mitochondria. Mitochondrial parameters were quantified using FiJi-based analysis tools. To quantify ER-plasma membrane (PM) interactions, PM and adjacent ER membranes were manually labeled in each image, at a single pixel thickness. The total length of ER segments adjacent to the PM was divided by the total PM length to obtain the interaction ratio.

### Focused ion beam-scanning electron microscopy (FIB-SEM) analysis

#### Sample preparation and imaging

For FIB-SEM imaging, samples were prepared using the rOTO protocol as previously described^25^ and embedded in Epon resin. A total of four samples were used, including one BAT sample from STIM1/2^flox^ mice maintained at room temperature (imaged volume: 1029 μm^3^), two BAT samples from STIM1/2^flox^ mice exposed to cold (imaged volume: 2758 μm^3^ and 546 μm^3^) and one BAT sample from STIM1/2^AdpKO^ mice exposed to cold (imaged volume: 3163 μm^3^). Trimmed sample blocks were mounted onto aluminum stubs using silver paint (Ted Pella Inc.) and then sputter-coated with a palladium/gold (Pd/Au) layer using a Tousimis sputter coater on top of a Bio-Rad E5400 controller. FIB-SEM imaging was performed with a Zeiss Crossbeam 550 (Carl Zeiss Microsystems GmbH, Oberkochen, Germany). Samples were tilted at 54° perpendicular to ion beam. The FIB milling and SEM imaging of the target area were set up using Atlas 5 3D tomography (Carl Zeiss Microsystems GmbH, Oberkochen, Germany). Slices with a thickness of 6nm were milled from the target area using the 30 kV 300pA ion beam. Energy-selective Backscattered (ESB) images were collected at 1.5 kV 1nA, with a dwell time of 18 ns, image pixel size of 6nm, and tilt correction angle of 36°. The acquired images were aligned with the Slice Registration tool in Dragonfly 2022.2 (Comet Technologies Canada Inc., Canada).

#### FIB-SEM segmentation and data analysis

Ground truth labels were generated by manually annotating each class (ER, mitochondria, nucleus and lipid droplets) in Napari (v0.4.18). Tunable 2D-U-Net networks (DLSIA) were used to obtain rough predictions for each class^52^. These predictions were manually proofread and corrected in Napari. A block consisting of at least 250x250x250 voxels was used to train and fine-tune 3D-U-Net network models with Incasem^53^. Binary tiff masks (for ER and nucleus) and instance-based segmentation (for mitochondria and LD) were generated, to assign a unique identifier to each organelle. The false positive connections between different mitochondria were improved by the instance-based segmentation. Additional proofreading and manual corrections were performed in Zeiss Arivis Pro. Objects, images, videos and quantifications from each class were generated using Zeiss Arivis Pro (v4.2).

#### Mitochondria quantification

Mitochondria complexity index was calculated by the following formula MCI^2^ = SA^3^ / (16 p^2^V^2^) where SA is surface area in μm^2^ and V is volume in μm^3^, as previously described^54^.

#### Cortical ER and remaining ER network analysis

In **Fig. 1E**, the overall distribution and organization of the ER network was analyzed. To define the cortical ER, the cell volume was computationally eroded by 20 pixels (120 nm) in all directions and subtracted from the original cell volume, yielding a cortical shell. ER voxels within this cortical shell volume were classified as cortical ER (red in **Fig. 1E**), while the remaining ER voxels within the non-cortical region were classified as remaining ER (purple in **Fig. 1E**).

#### ER distance map from the plasma membrane

For the analysis in **Fig. 1F**, manually drawn plasma membrane and ER traces were used. A pixel-wise distance map was generated in Fiji for each image, where each ER pixel was assigned, a value based on its proximity to the plasma membrane: pixels closest to the membrane were assigned a value of 0 (0 nm), and the farthest ER pixels were assigned a value of 255 (1530 nm).

#### ER-plasma membrane interaction quantification

For the analysis in **Fig. 1G**, the plasma membrane and ER were manually traced in 469 images from BAT from mice at room temperature dataset and 440 consecutive images from BAT of cold-exposed mice dataset. 3D reconstructions were generated using Zeiss Arivis Pro software. To identify ER-PM contact sites, the ER traces were computationally expanded by three pixels (18 nm) in each plane, and pixels intersecting with the plasma membrane were defined as ER-PM contact sites.

#### ER-mitochondria interaction quantification

For the analysis in **Fig. 1I,** a single-voxel-thick representation of the mitochondrial outer surface was generated by plane-wise single pixel erosion of the mitochondria volume followed by subtraction from the original volume. ER-annotated voxels were then computationally expanded by three pixels (18 nm) in each plane. Voxels overlapping between the expanded ER volume and the mitochondrial outer surface were defined as the “interaction surface.” The percentage of mitochondrial surface in contact with the ER was calculated by dividing the interaction surface by the total single-voxel-thick mitochondrial outer surface area.

### Differentiated brown adipocytes

Brown preadipocyte cells (WT1 cells) were kindly donated by Alexander Bartelt (Technical University of Munich, Munich, Germany). Cells were cultured in Dulbecco’s modified Eagle’s medium (DMEM) with 4.5 mM glucose (Gibco) supplemented with 10% Cosmic calf serum (CCS, Hyclone) and 1% penicillin/streptomycin (Gibco) at 37 °C, 5% CO2. Once the cells reached 100% confluence (2 to 3 days after seeding), the medium was changed to DMEM containing 10% fetal bovine serum (FBS, R&D Systems) with 1% penicillin/streptomycin. After 2 days (day 0), adipocyte differentiation was induced by the addition of 500 μM 3-isobutyl-1-methylxanthine, insulin (5 μg/ml), 10 μM dexamethasone, and 10 μM rosiglitazone to the culture media. On day 2, the medium was switched to DMEM with 10% FBS and 1% penicillin/streptomycin, insulin (5 μg/ml), and 10 μM rosiglitazone (maintenance medium) and was replaced every other day until day 8-10, when the adipocytes were fully differentiated and experiments could be performed.

### Primary adipocyte isolation, differentiation and culture

Neonatal pups (1-3 days old) were euthanized by decapitation, and interscapular BAT depots were aseptically dissected and transferred into a Petri dish containing ice cold PBS. For each isolation, BAT from 3 to 4 pups were pulled together, tissues were minced and transferred to a HEPES-buffered solution (pH 7.4) containing 123 mM NaCl, 5 mM KCl, 1.3 mM CaCl2, 5 mM glucose, 2% BSA (fatty acid free), 100 mM HEPES, and collagenase type II (Worthington). Minced tissues were pooled in 2.5 mL of the previously described HEPES-buffered solution per mouse and digested at 37°C on a shaker at 100 rpm for 40 minutes, with brief vortexing every 10 minutes. For every 10 mL of the digest, 5 mL of DMEM (w/ 1% PenStrep, no serum) was added to the digest. The digest was further disrupted with a 18G needle 8-10 times and 20-25 mL of DMEM (w/ 1% PenStrep, no serum) was added before being filtered through a 40-μm nylon filter into a 50-mL sterile tube. The tube was centrifuged for 10 min at 600g. The pellet was resuspended in DMEM and centrifuge once more at 600g for 10 min, to wash the cells. The resulting pellet was then resuspended in 2 mL/mouse of cell culture medium containing DMEM, 15% FBS, and 1% PenStrep. The cell suspension was seeded into a 10 cm plate. Two-day after isolation, the cells were washed with pre-warmed DMEM and replaced with cell culture medium. The medium was changed every 2 days until cells are 100% confluent. Differentiation was induced in the same manner as described above with WT1 cells for 5-7 days.

### Cell transfections

For the downregulation of STIM1/2 in WT1 brown adipocytes, cells were transfected with scrambled siRNA or siRNA against STIM1 and 2 (in combination) at day 10 of differentiation. Briefly, cells were trypsinized (0.05% trypsin), transferred into 15-ml conical tubes, and centrifuged for 10 min at 300 rpm at 4°C. The supernatant was removed, and the pellets were diluted in transfection medium (DMEM, 10% FBS, and no penicillin/streptomycin). The adipocytes were counted and seeded in 6- or 12-well plates. Transfection mixtures were prepared in Opti-MEM following the Lipofectamine RNAiMAX protocol and added to the culture medium.

Cells were incubated at 37 °C for 16 hours, after which the medium was replaced with maintenance medium. Cells were then incubated for an additional 20 hours (36 hours total post-transfection) prior to experimentation. Lipofectamine RNAiMAX (Thermo, # 13778030) and Opti-MEM medium (Thermo, #31985062); mouse siRNAs for ON-TARGET SMARTpool Non-targeting Control (catalog ID: D-001810-01-05, STIM1 (L-062376-00-0005), STIM2 (L-055069-01-0005), were from Dharmacon.

### MAPPER based ER-PM imaging

GFP-MAPPER was a gift from Dr. Jen Liou (Addgene plasmid # 117721)^29^. Lentivirus particles were generated by co-transfection of this plasmid with third-generation lentiviral packaging plasmids (pMDLg/pRRE and pMD2.G) into HEK293T cells. Media containing lentivirus particles was collected 72 hours after transfection, passed through a 40 µm filter, and then used to infect differentiated WT1 brown adipocytes. For infection, WT1 cells were seeded into 6-well imaging plates. Lentivirus particles were added in the presence of 10 μg/ml Polybrene. To increase infection rate, cells were centrifuged at 800g at 25°C for 90 min. Brown adipocytes expressing MAPPER were imaged 48 hours after infection with a Yokogawa CSU-X1 spinning disk confocal with a Zeiss AxioObserver Z1 microscope (Carl Zeiss Microscopy), with PlanApo 63x/1.4 and PlanApo 100x/1.4 objectives, equipped with an Omicron LaserAge Light Hub (488nm laser), Hamamatsu ImageEM X2 EMCCD camera, Okolab environmental system, and controlled by MetaMorph software, version 7.10.5.476 (Molecular Devices, LLC). To quantify GFP-positive puncta at the bottom plane of the cells, cell borders were manually drawn. Puncta were thresholded to generate binary images, and maxima were identified to mark individual puncta. The number of puncta was then normalized to the corresponding cell area to yield puncta density per cell.

### Total protein extraction and western blotting

Brown adipose tissue was collected from mice and any white adipose tissue surrounding BAT was carefully removed. Tissues were homogenized in cold lysis buffer containing 50mM Tris-HCl (pH 7.4), 2mM EGTA, 5mM EDTA, 30mM NaF, 10mM Na3VO4, 10mM Na4P2O7, 40mM glycerophosphate, 1 % NP-40, and 1% protease inhibitor cocktail. After ∼20 minutes of incubation on ice, the homogenates were centrifuged at 9,000 rpm for 15 minutes to pellet cell debris. The resulting supernatant was collected, and protein concentrations were determined using the BCA assay. Samples were diluted in 4x Laemmli buffer and boiled for 5 min. Protein lysates were subjected to SDS-polyacrylamide gel electrophoresis and transferred to PVDF membranes. Membranes were blocked in the presence of 3% BSA. All the immunoblots were incubated with primary antibody overnight at 4°C, washed in TBS-T at least 3-4 times for 5 mins followed by incubation with secondary antibody conjugated with horseradish peroxidase for 1-3 hours at room temperature. After incubation, membranes were washed with TBS-T and luminescence signal was visualized using the enhanced chemiluminescence system (Bio-Rad). Antibodies used were STIM1 (Cell Signaling, 4916), STIM2 (Cell Signaling, 4917), α-Tubulin (Proteintech, 66031-1-Ig), β-Tubulin (Cell Signaling, 2146S), GAPDH (Cell Signaling, 5174S), UCP1 (Cell Signaling, 14670); Junctophilin (JH2) (Proteintech, 51054-2-AP) ; pHSL (Cell Signaling, 4139) ; HSL (Cell Signaling, 4107) ; Oxphos cocktail (Abcam, ab110413); pDRP1 ser616 (Cell Signaling, 4494); DRP1 (Cell Signaling, 8570), anti-rabbit IgG (Cell Signaling, 7074S), and anti-mouse IgG (Cell Signaling, 7076S).

### Gene expression analysis

Brown adipose tissues were disrupted in TRIzol (Invitrogen) using TissueLyser (Qiagen). To obtain RNA, Trizol homogenates were mixed with chloroform vortexed thoroughly and centrifuged at 12000g for 20 min at 4°C. The top layer was transferred to another tube and mixed with 70% Ethanol. The RNA was extracted following the NucleoSpin RNA kit protocol (Macherey-Nagel, 740955.50) and eluted in RNAse free water. Complementary DNA (cDNA) was synthesized using iScript RT Supermix kit (Biorad). Quantitative real-time PCR reactions were performed in duplicates or triplicates on a LightCycler 480 II System (Roche) using SYBR green and custom primer sets, or primer sets based on the Harvard primer bank. Gene of interest cycle thresholds (Cts) were normalized to TBP (TATA box–binding protein) housekeeping levels or to Adiponectin. Expression levels were quantified using the ΔΔCt method. The sequences of the primers used for qPCR are listed in Table 1.

**Table 1:**
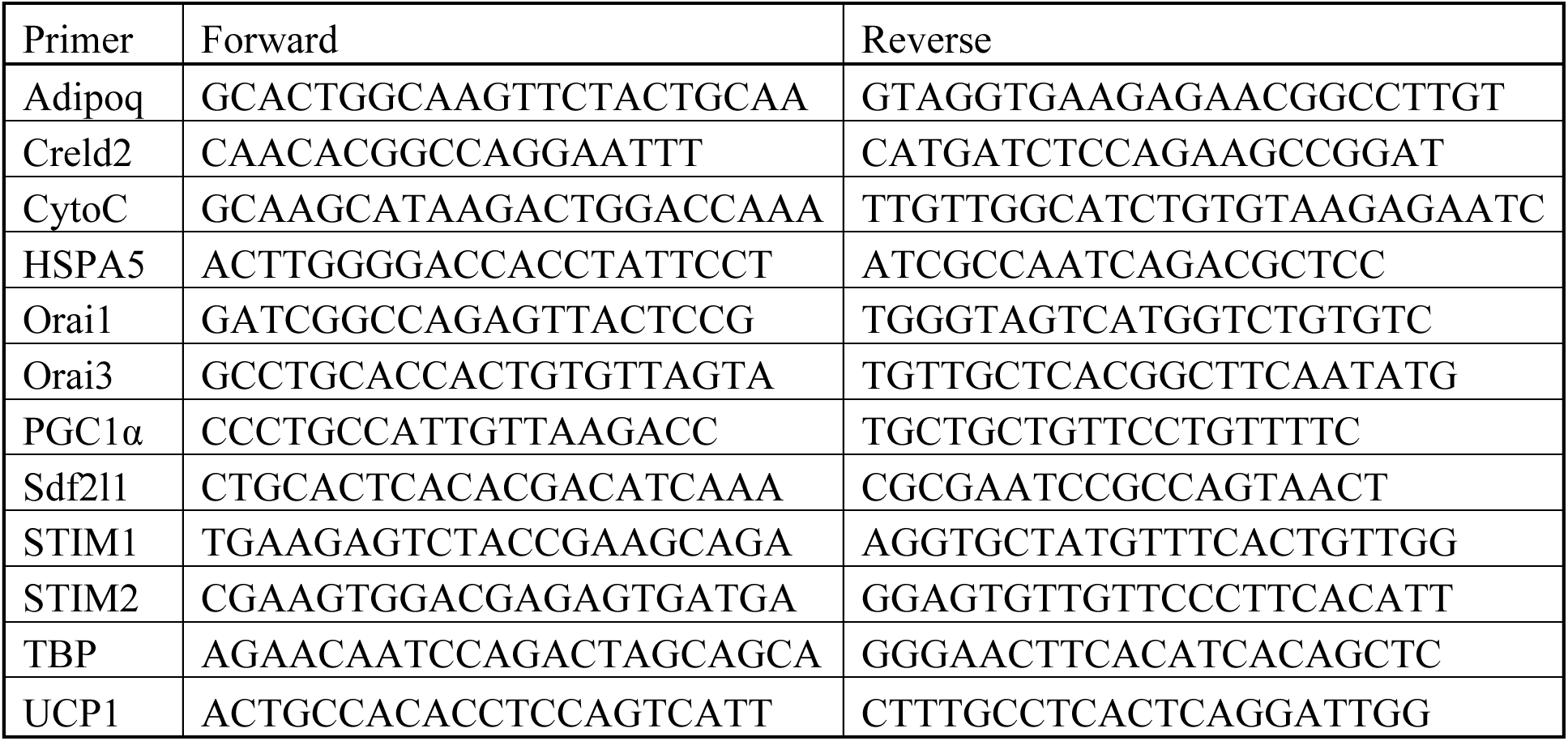
Sequence of primers used in qPCR analysis.

### Immunostaining

To assess the endogenous localization of STIM1 protein, differentiated brown adipocytes were incubated in calcium-free medium containing either 1µM thapsigargin, DMSO (vehicle control), or 1 or 10 µM norepinephrine, for the durations indicated in the figure legends. Cells were fixed with 4% paraformaldehyde for 10 minutes at room temperature (RT), followed by three washes in PBS. Permeabilization was then performed for 20 minutes at RT using 0.2% Triton X-100 in 2% BSA. STIM1 was stained with Rabbit anti-STIM1 (Cell Signaling, #4916) antibody overnight at 4°C, at a concentration of 1:200 in PBS. The next day, cells were washed 3x, including one long wash for more than 10 min. The cells were incubated with secondary antibody Alexa fluor 488 (Abcam, ab184819), diluted 1:1000 in PBS, at RT for 1hr in the dark. The cells were washed 3x, including one 10 min wash, and if needed, lipid droplets were labeled with 1 µL AUTODOT (Abcepta, SM1000a) per dish before imaging. Nucleus were stained with Hoechst. For the staining of mitochondria, differentiated primary brown adipocytes were fixed, permeabilized and blocked using the same protocol as described above. Mitochondria were stained with total OXPHOS Rodent Antibody Cocktail (Abcam, ab110413) at 1:200 dilution, nucleus were labeled with Hoechst. Cells were imaged with a Yokogawa CSU-X1 spinning disk confocal with a Zeiss AxioObserver Z1 microscope (Carl Zeiss Microscopy), with PlanApo 63x/1.4 and PlanApo 100x/1.4 objectives, equipped with an Omicron LaserAge Light Hub (488nm laser), Hamamatsu ImageEM X2 EMCCD camera, Okolab environmental system, and controlled by MetaMorph software, version 7.10.5.476 (Molecular Devices, LLC).

#### Analysis of STIM1 translocation to plasma membrane contact sites

To assess STIM1 translocation toward the plasma membrane, rectangular regions of interest (ROIs) of identical size were manually drawn over areas corresponding to the plasma membrane and adjacent cytosolic regions. The mean fluorescence intensity within each ROI was measured, and STIM1 translocation was quantified as the ratio of plasma membrane fluorescence intensity to adjacent cytosolic fluorescence intensity.

#### Analysis of Mitochondria shape

Mitochondria were segmented using MitoSegNet^55^ deep-learning model from fluorescent images in each cell of interest (**Extended Fig. 5C**). Mitochondria feature parameters were then quantified using FiJi.

### Calcium imaging

Cytosolic Ca^2+^: For the single cell cytosolic Ca^2+^ measurements presented in **Figures 2D** and **2F**, cells were loaded with 4μM Fluo4 in HBSS medium for ∼25 to 40 min at room temperature. Cells were then washed and kept in Ca^2+^ free medium containing 10mM Hepes, 150mM NaCl, 4mM KCl, 3mM MgCl2, 2mM EGTA, 10mM D-glucose, pH 7.4. Baseline signal was recorded. Cells were then treated with 1μM thapsigargin (**Fig. 2D**) or 10μM norepinephrine (**Fig. 2F**) in Ca^2+^ free medium. In the indicated time points cells were loaded with a medium containing 10 mM HEPES, 150mM NaCl, 4mM KCl, 1mM MgCl2, 2mM CaCl2, 10mM D-glucose, pH 7.4 and fluorescence intensity was recorded. For the single cell cytosolic Ca^2+^ measurements presented in **Figure 2G**, cells were loaded with Fluo4 as described above, washed and incubated in medium containing 10mM HEPES, 150mM NaCl, 4mM KCl, 1mM MgCl2, 2mM CaCl2, 10mM D-glucose, pH 7.4. Norepinephrine (10uM) was added to the cells at the indicated time point and changes in fluorescence were recorded. For all the experiments images were taken every 2 or 5 seconds with a 20x objective in a Yokagawa spinning disk confocal microscope equipped with an Omicron laser merge module (488 laser), Hamamatsu ImageEM X2 EMCCD camera. Metamorph was the software used for acquisition parameters, shutters, filter positions and focus control. Images were background corrected and analyzed with FiJi software. Average of traces of stimulant responsive cells are shown in the figures. Mitochondrial Ca^2+^: For the single cell mitochondrial Ca^2+^ uptake presented in **Fig. 2H**, ells were loaded with 4μM Rhodamine-2 (Rhod-2) in HBSS medium for ∼30 to 40 min at room temperature. Cells were washed and placed in a medium containing 10 mM HEPES, 150mM NaCl, 4mM KCl, 1mM MgCl2, 2mM CaCl2, 10mM D-glucose, pH 7.4 and fluorescence intensity was recorded. Cells were treated with 10μM norepinephrine in the indicated time point.

### Cold exposure challenge

For cold exposure experiments, mice were transferred from room temperature to a cold room maintained at 4°C and housed individually in cages with bedding and ad libitum access to food and water unless specified in the legend. Mice were exposed to cold from 6 to 24 hours depending on the experiment and the times are specified in the legend of the figures. For the experiments presented in **Figures 3B** and **3C** and **3F** cold exposure was performed in the absence of food. Mice were removed from the cold chamber if their body temperature dropped below 25°C, as approved in our protocol. Core body temperature was monitored every 30 minutes using a RET-3 Rectal Probe (Physitemp). To assess the BAT surface temperature (**Figure 3D**), mice were exposed to cold for 4-6 hours, briefly anesthetized with isoflurane, and the skin overlying the brown adipose depot was carefully removed. Each mouse was then placed on a flat surface, and thermographic images were captured using a FLIR E5 series infrared camera. The camera was positioned at a fixed distance from the surface to ensure consistent measurements across all animals.

### Indirect calorimetry in anesthetized mice

Mice were anesthetized by an intraperitoneal injection of ketamine/xylazine (300 mg/kg ketamine and 30 mg/kg xylazine in PBS) and placed in individual metabolic cages Comprehensive Lab Animal Monitoring System (CLAMS, Columbus Instruments) at room temperature. Data was collected from four mice at a time. After 8 min of baseline measurements, cages were opened and norepinephrine (Sigma-Aldrich, A9512, diluted in isotonic saline to a concentration of 0.1mg/ml), was subcutaneously injected to mice (injection volume: 200 ul per mouse) above the interscapular brown adipose tissue, resulting in 20 mg NE administered per mouse.

### Histological analysis

Brown adipose tissues were fixed in 10% zinc formalin for 24hours at room temperature and transferred to 70% ethanol for further storage. Tissues were processed, sectioned, and stained with hematoxylin and eosin at the UCSF Liver Center Pathology Core or at Histowiz, Inc. (Brooklyn, NY). Lipid droplet area was analyzed with a custom macro generated in FiJi. Briefly, a Random-Forest-based machine-learning approach was used to segment the pixels occupied by lipid droplets (white colored regions). Binary masks were created and morphological features and numbers were quantified with ‘Analyze Particles’ function in FiJi.

### RNA sequencing and analysis

RNA from brown adipose tissue of mice maintained at room temperature or exposed to cold for 6 hours were extracted as described above (in qPCR section). RNA samples were submitted to University of California Davis Genetic Core. Barcoded 3’Tag-Seq libraries were prepared using the QuantSeq FWD kit (Lexogen, Vienna, Austria) for multiplexed sequencing according to the recommendations of the manufacturer. The fragment size distribution of the libraries was verified via micro-capillary gel electrophoresis on a Bioanalyzer 2100 (Agilent, Santa Clara, CA). The libraries were quantified by fluorometry on a Qubit instrument (LifeTechnologies, Carlsbad, CA) and pooled in equimolar ratios. Thirteen libraries were sequenced on one lane of an Aviti sequencer (Element Biosciences, San Diego, CA) with single-end 100 bp reads. The sequencing generated more than 4 million reads per library.

Single-end 100bp AVITI reads were subjected to quality control. Six base pair unique molecular index (UMI) was extracted using umi-tools/v1.1.2. Adapter sequences and low-quality bases (q < 30) were removed using trim_galore/v0.6.7. Reads that are less than 30bp in length were discarded. Reads that have passed the quality control were aligned to the mouse reference genome (GRCm39) and GENCODE annotation version 29 using STAR/v2.7.10b. PCR duplicates were removed using umi-tools based on the UMI information generated earlier in the workflow. Following the removal of PCR duplicates, gene expression was quantified using htseq-count function from HTSeq/v0.12.3 package. Raw counts were filtered to remove lowly expressed genes using filterByExpr function from edgeR and followed by TMM normalization. Differential expression analysis was carried out using limma+voom. Multiple test correction used Benjamini-Hochberg metho. Ingenuity Pathway analysis was used to define enriched pathways. Analysis of ATF6 gene targets shown in **Figure 4G**, was performed as previously described^40^. Briefly, ATF6- and XBP1s-selective target genes were identified from transcriptional profiling of HEK293DAX cells following stress-independent activation using TMP-inducible DHFR-ATF6 and/or doxycycline-inducible XBP1s. PERK-selective target genes were obtained from^56^. For this analysis, ATF6-, XBP1s-, or PERK-selective genes that were induced by more than 1.5-fold in thapsigargin (Tg)-treated samples were included.

### Glucose and insulin tolerance tests

For the oral glucose tolerance test, mice were administered glucose (Teknova, G9005) by gavage after 16hours overnight fasting (0.5 to 1.0 g kg^−1^), and blood glucose levels were measured throughout 120 min as indicated. Serum Insulin levels were measured using the Ultra-Sensitive Mouse Insulin ELISA kit (Crystal Chem, 90080). For insulin tolerance test, insulin (Humulin R, Eli Lilly) was injected intraperitoneally (0.5 to 1.0 U/kg^−1^) after 6hours of food withdrawal and blood glucose levels were measured throughout 120 min as indicated.

### Seahorse Respirometry in isolated mitochondria

#### Mitochondria isolation

Mitochondria were purified from brown adipose tissue (BAT) as described previously^57^ with modifications. BAT from three to four 6-8-week-old mice per experimental group was minced in a glass Petri dish containing 2 ml of sucrose-Tris-EGTA (STE) buffer (250 mM sucrose, 5 mM Tris-HCl, 2 mM EGTA, pH 7.4) and transferred to a 10 ml glass homogenizer with a Teflon pestle containing 5 ml STE supplemented with 1% (wt/vol) fatty acid–free BSA. The tissue was manually homogenized (up to six strokes), the volume was adjusted to 25 ml, and the suspension was centrifuged at 8,500 g for 10 min. The fat layer was discarded, and the tube walls were wiped clean to remove residual fat. The pellet was resuspended in 1 ml of BSA-free STE, further homogenized in a glass–glass homogenizer, and transferred to a clean tube. The volume was brought up to 25 ml, and the sample was centrifuged at 800g for 5 min. The pellet was discarded, and the supernatant was collected and centrifuged at 8,500g for 10 min. The pellet was washed in BSA free media and resuspended in 150μl STE. All steps were performed at 4 °C. Protein concentration was determined by the BCA assay (Pierce).

#### Seahorse Respiration assay

5 μg mitochondrial protein were added per well in a media containing 50 mM KCl, 10 mM TES, 1 mM EGTA, 0.4% (wt/vol) FA-free BSA, 1 mM KH2PO4, 2 mM MgCl2 and 0.46 mM CaCl2 in the presence of 1 mM ADP and 3 mM GDP. The X24 plate was centrifuged at 2,000g for 15 min. The oxygen consumption rates were monitored in a Seahorse XF24 instrument. Baseline measurements (state 2) were performed, followed by injection of 20mM Pyruvate and 20mM Malate in port A and then 2 μg/ml oligomycin in port B. FCCP was titrated to 9 μM and injected in port C. Mitochondrial oxygen consumption was inhibited by the injection of 2μM rotenone and 2μM antimycin A in port D. The standard protocol was 40 sec mix, 2.5 min measure.

### Statistics and reproducibility

Statistical significance was assessed using GraphPad Prism version 9, by the unpaired Student’s t-test (two-tailed) or one-way or two-way ANOVA analysis as indicated in figure legends. P values are indicated in the figure legends. For in vitro studies a minimum of three biological replicates were used for each experiment.

## Data availability

Videos associated with this manuscript can be found in:

The uncropped raw versions of the western blots generated in the current study are provided as supplementary figures. Source data for all graphs are provided with this manuscript.

## Acknowledgements

We thank Luke Wiseman for his insight with the analysis of UPR genes show in **Figure 4G**. We thank all members of the Arruda laboratory for their continued support and encouragement. This project is supported by the Biohub, San Francisco grant to A.P.A; the NIH Pilot & Feasibility award P30DK098722 to G.P. and A.P.A. G.P. is supported by Burroughs Wellcome Fund. The sequencing was carried out by the DNA Technologies and Expression Analysis Core at the UC Davis Genome Center, supported by NIH Shared Instrumentation Grant 1S10OD010786-01. FIB-SEM data collected using the Zeiss XB 550 was supported by NIH S10 grant S10OD030258.

## Contributions

A.P.A designed and supervised the project, performed in vitro and in vivo experiments, analyzed the data, prepared the figures and wrote the manuscript. J.Z. designed experiments, performed most of the in vitro and in vivo experiments, analyzed the data and wrote the manuscript. L.L.A, C.D., S.H., K.R., S.H., K.L, T.K., V.X., G.P. assisted with in vivo and in vitro experiments. G.P. performed FIB-SEM preparation and data analysis, designed and supervised both in vitro and in vivo experiments, and analyzed the resulting data. G.P. also contributed intellectually to the development and progression of the project throughout. M.K., D.M.J. performed the image acquisition of the FIB-SEM data. S.B.W helped with RNA seq analysis. All authors read, edited, and contributed to the manuscript.

## Extended Figures

**Extended Figure 1.**
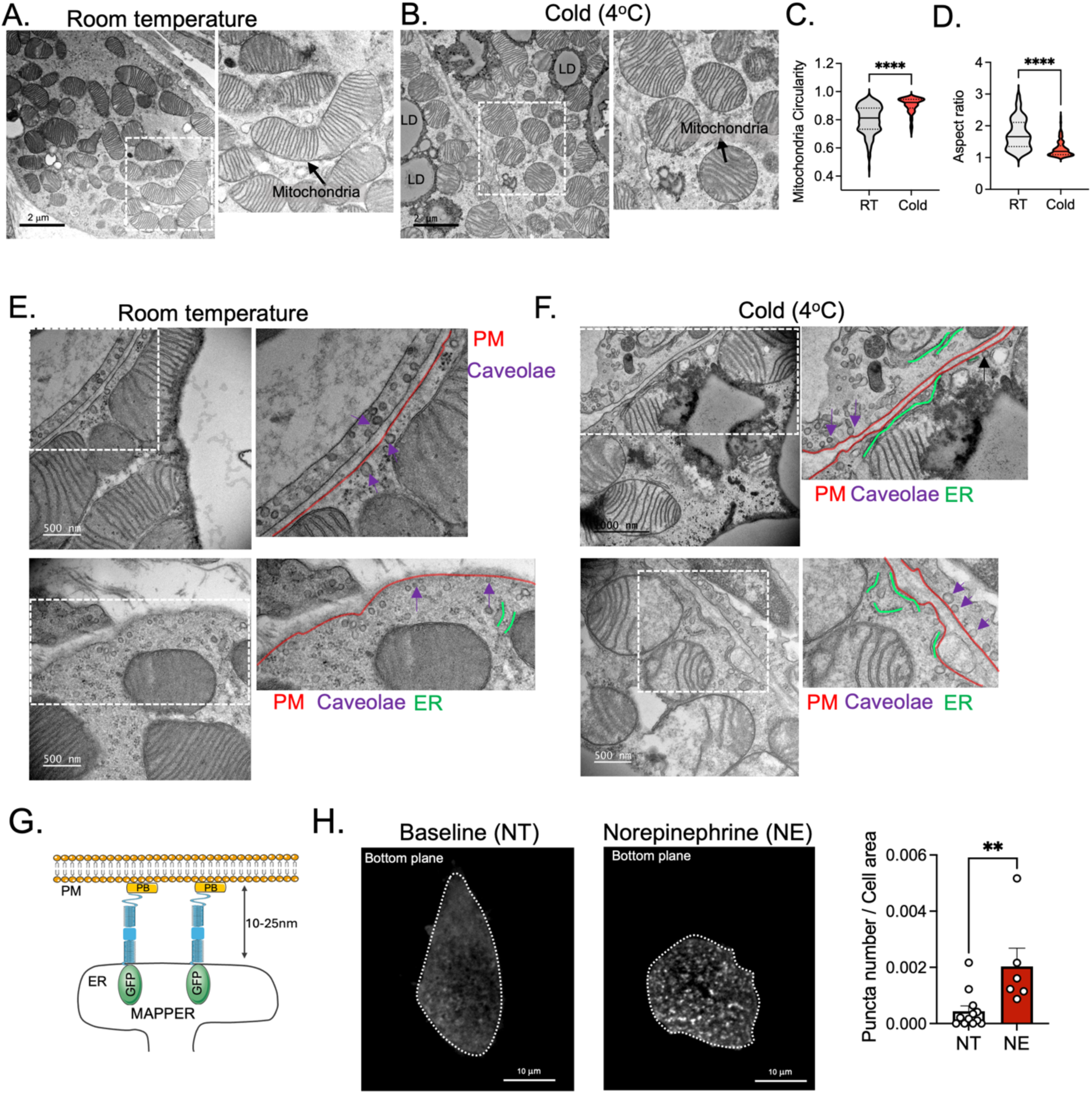
Cold exposure induces ER-PM contacts in brown adipocytes. (A, B) Representative transmission electron microscopy (TEM) images of brown adipose tissue from C57B6/J wild-type mice housed at room temperature (RT) (A) or exposed to cold (4°C) for 12 hrs (B). (C, D) Quantification of mitochondria circularity (C) and aspect ratio (D) from images shown in A and B. RT, n= 198 mitochondria in 13 images from 3 different mice. Cold n= 202 mitochondria in 9 images from 3 different mice. Unpaired Student’s t-test *** p<0.001. (E, F) Representative TEM images from cortical areas of brown adipocytes within BAT tissue from mice housed at RT (E) or exposed to cold (4°C) for 12 hrs (F). Plasma membrane is outlined in red, caveolae is indicated with purple arrows and ER is outlined in yellow. (G) Cartoon depicting the MAPPER construct. (H) Left: Representative confocal images of bottom-sections of differentiated brown adipocytes transiently expressing MAPPER construct. Cells were treated with 10μM of norepinephrine (NE) for 30 minutes. Right: Quantification of MAPPER puncta signal for images shown in the left. n=12 cells NT, n= 6 cells NE, Unpaired Student’s t-test ****p<0.0001. Representative of 2 independent experiments. All the bar graphs are represented as mean ± s.e.m.

**Extended Figure 2.**
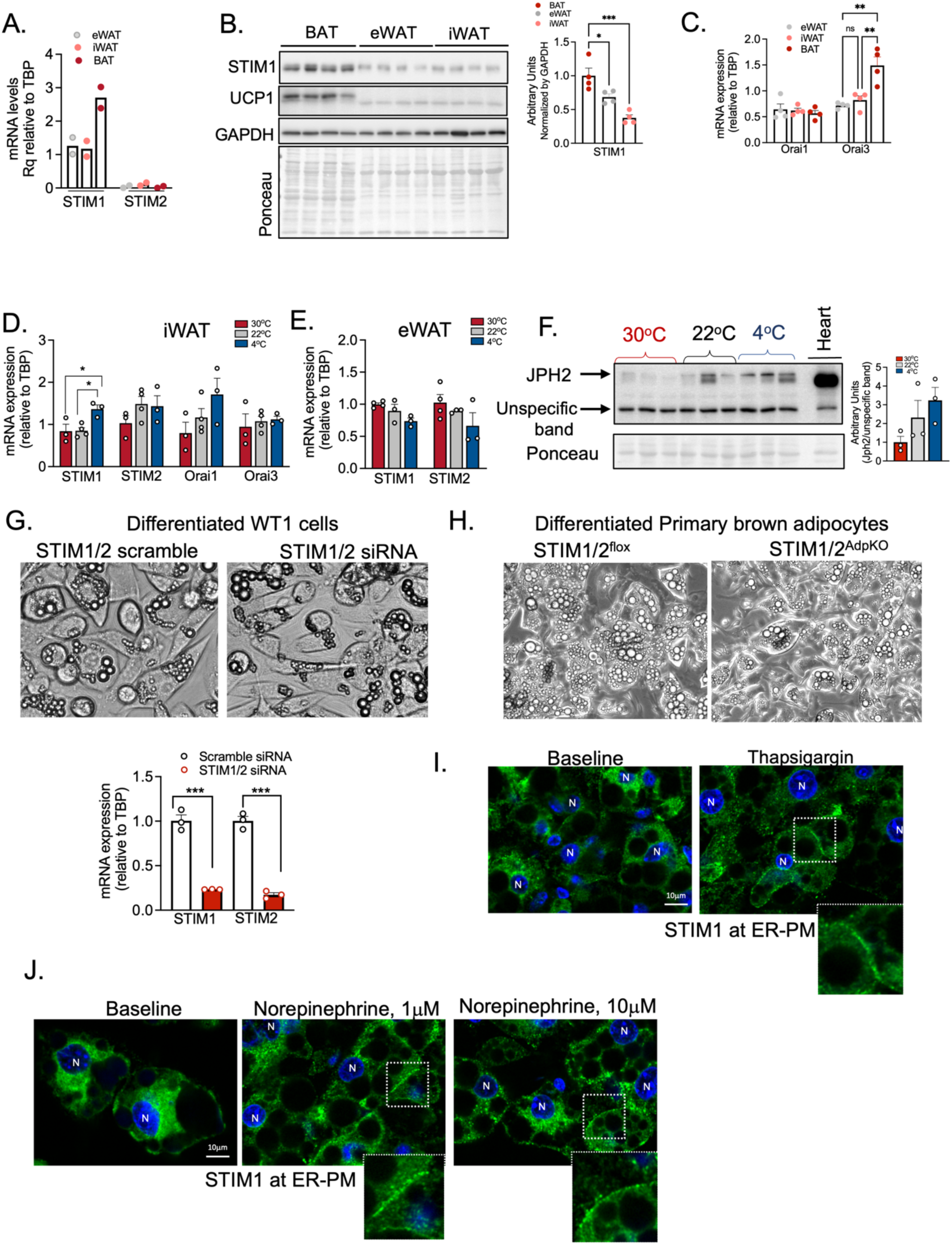
STIM and Orai expression levels and activity in adipocytes. (A) qPCR analysis of relative mRNA levels of indicated genes in BAT, epididymal white adipose tissue (eWAT) and inguinal white adipose tissue (iWAT). n= 2 mice per group (B) Left: Immunoblotting analysis of indicated proteins in total lysates from BAT, eWAT and iWAT from wild-type mice housed at RT. Right: Quantification analysis of the images shown in the left. n= 4 mice per group; One-way ANOVA, p< *0.05, *** p< <0.001. (C) qPCR analysis of relative mRNA levels of indicated genes in BAT, eWAT and iWAT. n= 4 per group; One-way ANOVA **p<0.005. (D, E) qPCR analysis of mRNA levels of indicated genes in iWAT (D) and eWAT (E) from mice exposed to 30°C, 22°C, and 4°C for 24hrs. In D, n= 3 (30°C); n=4 (22°C); n=3 (4°C); One-way ANOVA, * p<0.03. In E n= 4 (30°C); n= 4 (22°C); n=3 (4°C). (F) Left: Immunoblotting analysis of indicated protein in total lysates from BAT of mice housed at 30°C (thermoneutrality), 22°C (RT) or exposed to 4°C for 24 hrs. Right: Quantification analysis of images shown in the left. n= 3 per group (G) Upper panel: Representative bright field images of differentiated brown adipocytes (WT1 cells) control (scramble siRNA) or treated with STIM1/2 siRNA. Bottom panel: qPCR analysis of mRNA levels of indicated genes. n=3 per group. Unpaired Student’s t-test ***p<0.001. (H) Representative bright field images of differentiated primary brown adipocytes (after 7 days differentiation) from STIM1/2^flox^ and STIM1/2^AdpKO^. (I, J) Immunostaining of STIM1 (green) in differentiated brown adipocytes in baseline and in cells treated with 1μM thapsigargin for 30 min in Ca^2+^ free media (I) or 1μM or 10 μM Norepinephrine for 30 min in Ca^2+^ free media (J). Nucleus are stained in blue.

**Extended Figure 3.**
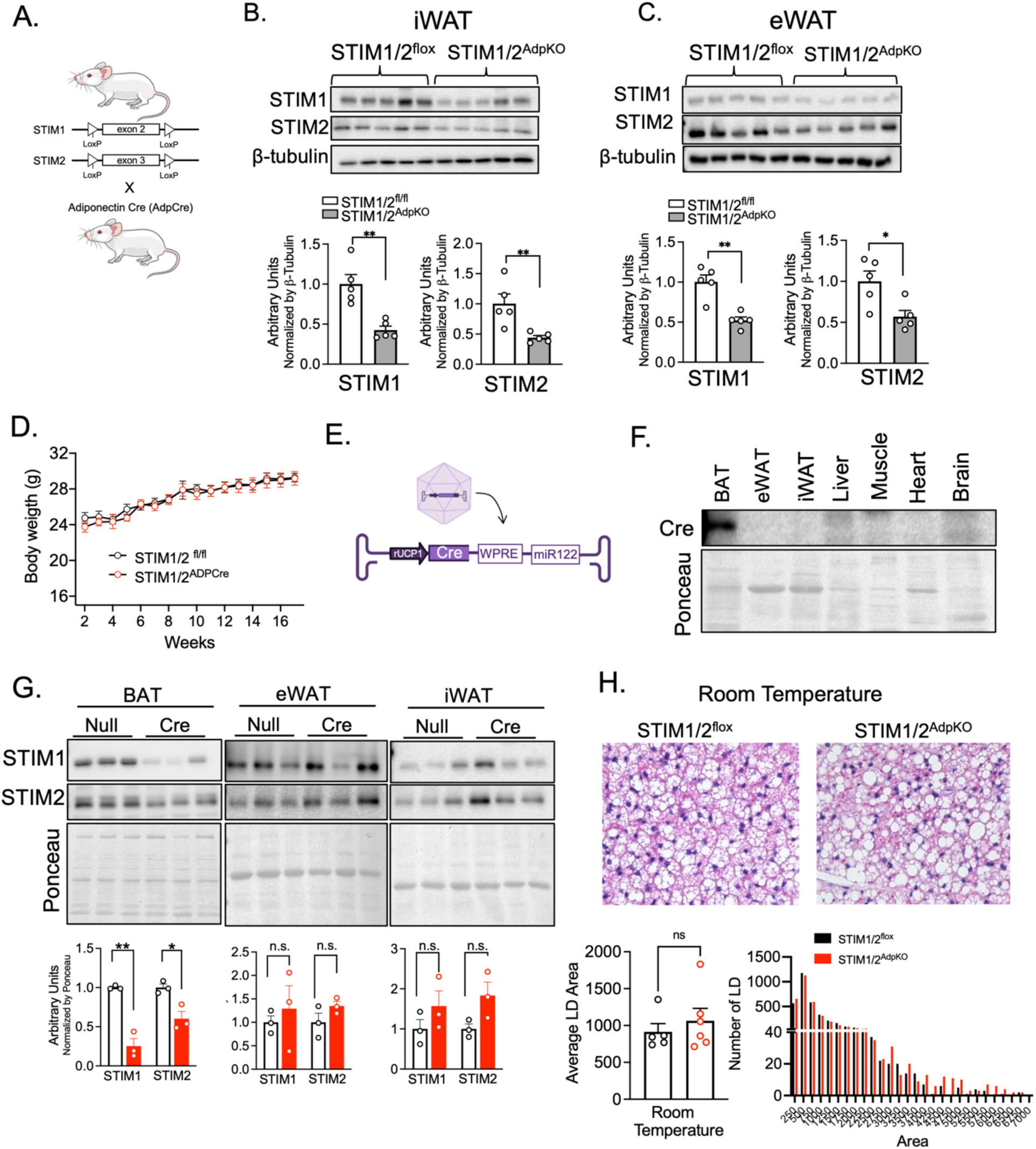
(A) Schematic depicting the genetic strategy used to generate mice with adipose tissue specific deletion of STIM1/2 (STIM1/2^AdpKO^). AdpCre- adiponectin Cre. (B) Upper panel: Immunoblotting analysis of indicated proteins in total lysates from iWAT of STIM1/2^flox^ and STIM1/2^AdpCre^ mice. Bottom panel: Quantification analysis of images shown in upper panel; n= 5 mice per group. Unpaired Student’s t-test, **p< 0.01. (C) Upper panel: Immunoblotting analysis of indicated proteins in total lysates from eWAT of STIM1/2^flox^ and STIM1/2^AdpKO^ mice. Bottom panel: Quantification analysis of images shown in upper panel; n= 5 mice per group. Unpaired Student’s t-test, * p<0.05, **p<0.002. (D) Body weight gain curves of STIM1/2^flox^ and STIM1/2^AdpKO^ mice fed a chow diet. n = 6 mice per group. (E) Schematic depicting the strategy used to generate mice with brown adipose tissue specific deletion of STIM1/2. (F) Immunoblotting analysis of Cre recombinase in the indicated tissues from mice injected with AAV-Rec2-UPC1-Cre-miR122. (G) Upper panel: Immunoblotting analysis of indicated proteins in in total lysates from BAT, eWAT and iWAT. Bottom panel: Quantification analysis of images shown in upper panel; n= 3 mice per group. Unpaired Student’s t-test, **p< 0.01, *p<0.05. (H) Top: Representative H&E-stained sections of BAT from STIM1/2^flox^ and STIM1/2^AdpKO^ mice housed at RT. Bottom: Quantification analysis of area of LD and frequence distribution.

**Extended Figure 4.**
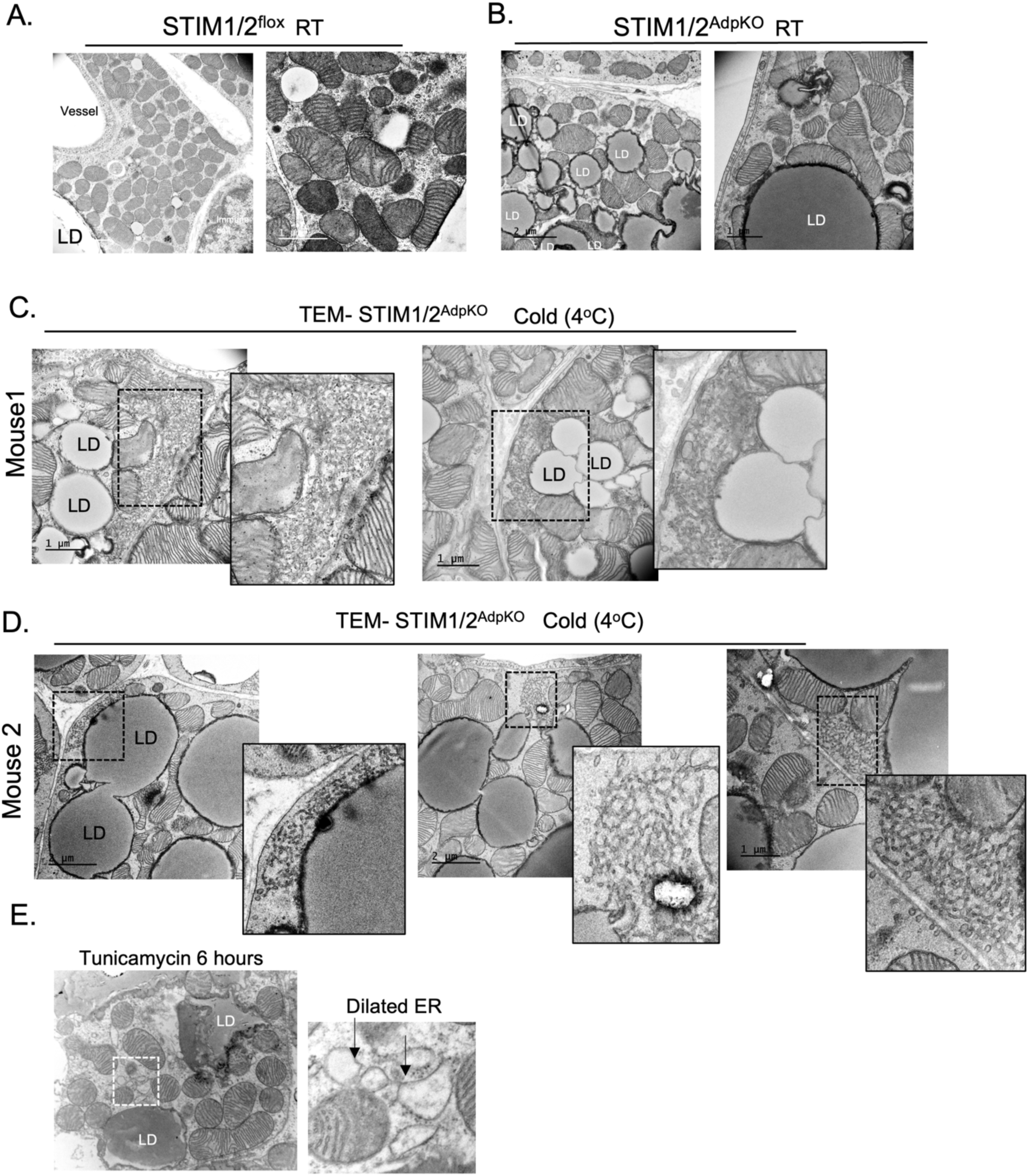
(A, B) Representative TEM images of BAT from STIM1/2^flox^ (A) and STIM1/2^AdpKO^ (B) mice maintained at RT. (C, D) TEM images of BAT from STIM1/2^AdpKO^ exposed to cold highlighting the aggregated membranes from two different mice. (E) Representative TEM image of BAT from mice treated with tunicamycin (0.5mg/kg) for 6 hrs.

**Extended Figure 5.**
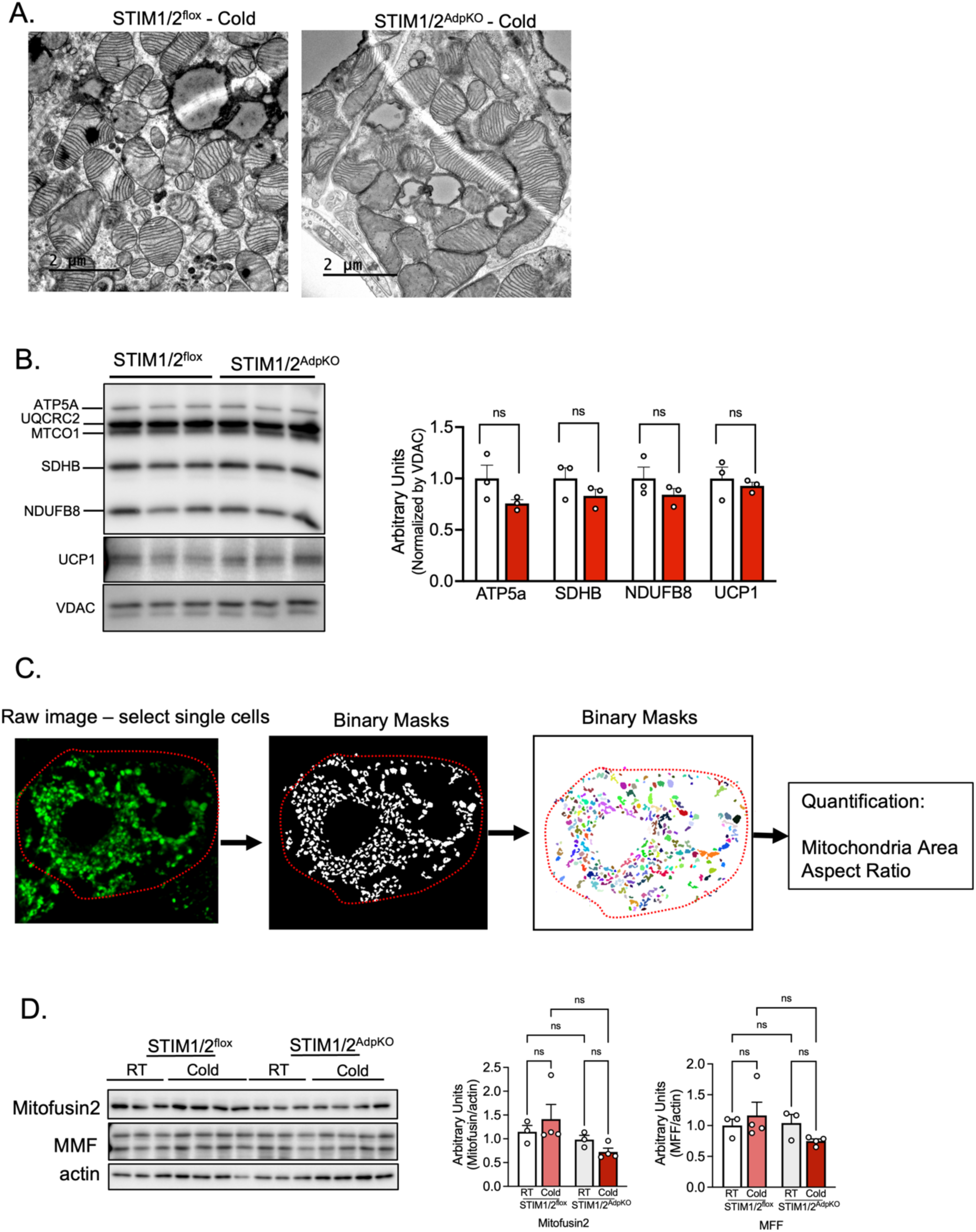
(A) Representative TEM images of BAT from STIM1/2^flox^ and STIM1/2^AdpKO^ exposed to cold (4°C). (B) Left: Immunoblotting analysis of indicated proteins in isolated mitochondria from BAT of STIM1/2^flox^ and STIM1/2^AdpCre^ mice exposed to cold for 12hrs. Right: Quantification analysis of images shown in left panel; n= 3 mice per group. (C) Quantification strategy for images shown in Fig. 5F. (D) Left: Immunoblotting analysis of indicated proteins in total lysates of BAT from STIM1/2^flox^ and STIM1/2 ^AdpKO^ mice maintained at RT or exposed to cold (4°C) for 2hrs. Right: Quantification analysis of images shown in the upper panel. n=3 mice per group for RT conditions and n=4 mice per group for cold conditions. One-way ANOVA with Tukey’s multiple comparisons test.

**Extended Figure 6.**
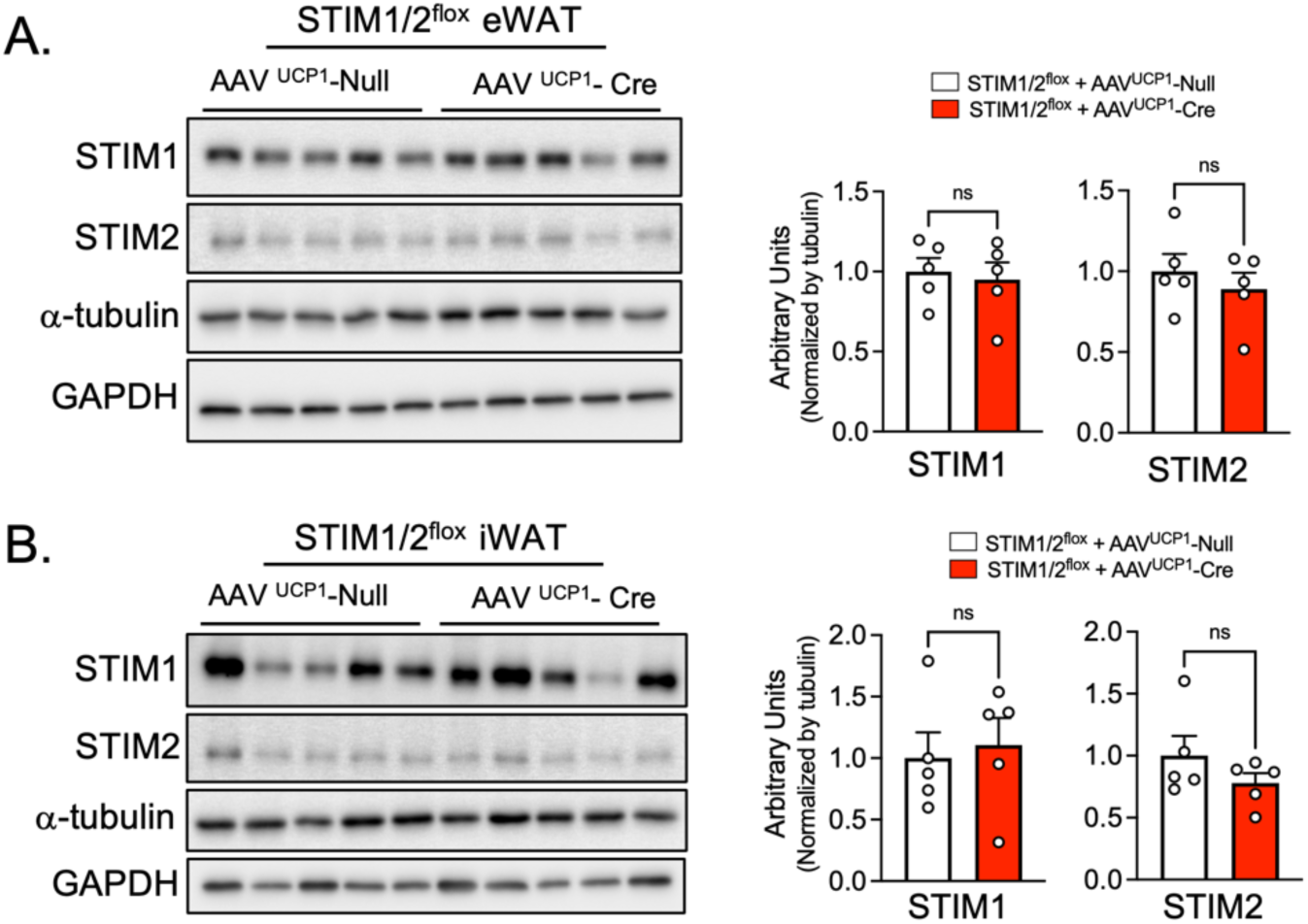
(A) Left panel: Immunoblotting analysis of indicated proteins in total lysates of eWAT of mice fed HFD infected with AAV-Rec2-UCP1-Null or AAV-Rec2-UCP1-Cre. Right panel: Quantification analysis of the images shown in left panel; n= 5 per group. (B) Left panel: Immunoblotting analysis of indicated proteins in total lysates of iWAT of mice fed HFD infected with AAV-Rec2-UCP1-Null or AAV-Rec2-UCP1-Cre. Right panel: Quantification analysis of the images shown in left panel; n= 5 per group.

**Extended Figure 7.**
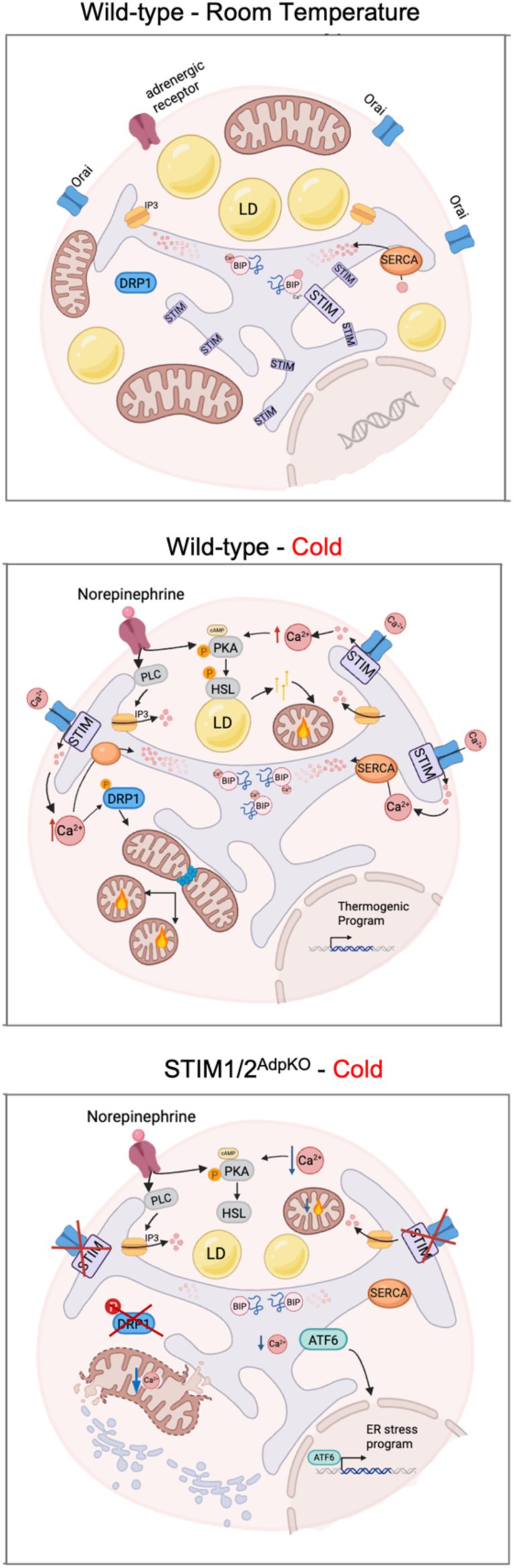
Model of the proposed mechanism of STIM1/2-mediated SOCE regulation of brown adipocyte adaptation to cold.

## Video Legends

**Video 1 –** 3D reconstruction of FIB-SEM data of BAT derived from mice housed at room temperature showing the ER, mitochondria, lipid droplets and nuclei.

**Video 2 –** 3D reconstruction of ER of BAT from mice housed at room temperature highlighting cortical ER at a distance of 120nm of the plasma membrane in red.

**Video 3 –** 3D reconstruction of ER of BAT from mice exposed to cold (4°C) for 12hrs highlighting cortical ER at a distance of 120nm of the plasma membrane in red.

**Video 4 –**3D reconstruction of FIB-SEM images from BAT of mice maintained at room temperature, highlighting regions of the plasma membrane where the endoplasmic reticulum (ER) is in close proximity (within 18 nm).

**Video 5 –** 3D reconstruction of FIB-SEM images from BAT of mice exposed to cold for 12hrs, highlighting regions of the plasma membrane where the endoplasmic reticulum (ER) is in close proximity (within 18 nm).

**Video 6 –** 3D reconstruction of FIB-SEM images reveals aggregated membranes accumulating in BAT tissue from STIM1/2-deficient mice following 12 hours of cold exposure.

**Video 7 –** 3D reconstruction of mitochondria morphology of BAT from STIM1/2 ^flox^ control mice exposed to cold (4°C) for 12hrs

**Video 8 –** 3D reconstruction of mitochondria morphology of BAT from STIM1/2 ^AdpKO^ control mice exposed to cold (4°C) for 12hrs

## References

1. Cannon, B., and Nedergaard, J. (2004). Brown Adipose Tissue: Function and Physiological Significance. Physiol. Rev. 84, 277–359. 10.1152/PHYSREV.00015.2003.

2. Cohen, P., and Kajimura, S. (2021). The cellular and functional complexity of thermogenic fat. Nat. Rev. Mol. Cell Biol. 22, 393–409. 10.1038/s41580-021-00350-0.

3. Sharma, A.K., Khandelwal, R., Zurkovic, J., Long, F., Wang, T., Dewal, R.S., Wu, C., Ghosh, A., Manuel, K., Othman, A., et al. (2026). DGAT-driven futile lipid cycling has a pronounced, yet concealed, thermogenic function. Cell Metab. 10.1016/J.CMET.2025.12.009.

4. Liu, X., He, A., Lu, D., Hu, D., Tan, M., Abere, A., Goodarzi, P., Ahmad, B., Kleiboeker, B., Finck, B.N., et al. (2025). Peroxisomal metabolism of branched fatty acids regulates energy homeostasis. Nature 646, 1223–1231. 10.1038/s41586-025-09517-7.

5. Rahbani, J.F., Roesler, A., Hussain, M.F., Samborska, B., Dykstra, C.B., Tsai, L., Jedrychowski, M.P., Vergnes, L., Reue, K., Spiegelman, B.M., et al. (2021). Creatine kinase B controls futile creatine cycling in thermogenic fat. Nature 590, 480–485. 10.1038/S41586-021-03221-Y.

6. Yoneshiro, T., Wang, Q., Tajima, K., Matsushita, M., Maki, H., Igarashi, K., Dai, Z., White, P.J., McGarrah, R.W., Ilkayeva, O.R., et al. (2019). BCAA catabolism in brown fat controls energy homeostasis through SLC25A44. Nature 572, 614–619. 10.1038/s41586-019-1503-x.

7. Bartelt, A., Bruns, O.T., Reimer, R., Hohenberg, H., Ittrich, H., Peldschus, K., Kaul, M.G., Tromsdorf, U.I., Weller, H., Waurisch, C., et al. (2011). Brown adipose tissue activity controls triglyceride clearance. Nat. Med. 17, 200–206. 10.1038/nm.2297.

8. Villarroya, F., Cereijo, R., Villarroya, J., and Giralt, M. (2017). Brown adipose tissue as a secretory organ. Nat. Rev. Endocrinol. 13, 26–35. 10.1038/nrendo.2016.136.

9. Müller, T.D., Blüher, M., Tschöp, M.H., and DiMarchi, R.D. (2022). Anti-obesity drug discovery: advances and challenges. Nat. Rev. Drug Discov. 21, 201–223. 10.1038/s41573-021-00337-8.

10. Cypess, A.M., and Kahn, C.R. (2010). Brown fat as a therapy for obesity and diabetes. Curr. Opin. Endocrinol. Diabetes Obes. 17, 143–149. 10.1097/MED.0B013E328337A81F.

11. Li, H., Li, J., Song, C., Yang, H., Luo, Q., and Chen, M. (2024). Brown adipose tissue: a potential target for aging interventions and healthy longevity. Biogerontology 25, 1011–1024. 10.1007/s10522-024-10137-3.

12. Chouchani, E.T., Kazak, L., and Spiegelman, B.M. (2019). New Advances in Adaptive Thermogenesis: UCP1 and Beyond. Cell Metab. 29, 27–37. 10.1016/j.cmet.2018.11.002.

13. Bartelt, A., Widenmaier, S.B., Schlein, C., Johann, K., Goncalves, R.L.S., Eguchi, K., Fischer, A.W., Parlakgül, G., Snyder, N.A., Nguyen, T.B., et al. (2018). Brown adipose tissue thermogenic adaptation requires Nrf1-mediated proteasomal activity. Nat. Med. 24, 292–303. 10.1038/nm.4481.

14. Zhou, Z., Torres, M., Sha, H., Halbrook, C.J., van den Bergh, F., Reinert, R.B., Yamada, T., Wang, S., Luo, Y., Hunter, A.H., et al. (2020). Endoplasmic reticulum-associated degradation regulates mitochondrial dynamics in brown adipocytes. Science. 2020 Apr 3;368(6486):54-60. doi: 10.1126/science.aay2494.

15. Latorre-Muro, P., O’Malley, K.E., Bennett, C.F., Perry, E.A., Balsa, E., Tavares, C.D.J., Jedrychowski, M., Gygi, S.P., and Puigserver, P. (2021). A cold-stress-inducible PERK/OGT axis controls TOM70-assisted mitochondrial protein import and cristae formation. Cell Metab. 33, 598–614.e7. 10.1016/j.cmet.2021.01.013.

16. De Meis, L., Arruda, A.P., Da Costa, R.M., and Benchimol, M. (2006). Identification of a Ca2+-ATPase in brown adipose tissue mitochondria: Regulation of thermogenesis by ATP and Ca2+. Journal of Biological Chemistry 281, 16384–16390. 10.1074/JBC.M600678200.

17. Auger, C., Li, M., Fujimoto, M., Ikeda, K., Yook, J.S., O’Leary, T.R., Caycedo, M.P.H., Xiaohan, C., Oikawa, S., Verkerke, A.R.P., et al. (2025). Identification of a molecular resistor that controls UCP1-independent Ca2+ cycling thermogenesis in adipose tissue. Cell Metab. 37, 1311–1325.e9. 10.1016/j.cmet.2025.03.009.

18. Wu H, Carvalho P, Voeltz GK. Here, there, and everywhere: The importance of ER membrane contact sites. Science. 2018 Aug 3;361(6401):eaan5835. doi: 10.1126/science.aan5835.

19. Arruda, A.P., and Parlakgül, G. (2023). Endoplasmic Reticulum Architecture and Inter-Organelle Communication in Metabolic Health and Disease. Cold Spring Harb. Perspect. Biol. 15. 10.1101/cshperspect.a041261.

20. Parlakgül, G., Pang, S., Artico, L.L., Min, N., Cagampan, E., Villa, R., Goncalves, R.L.S., Lee, G.Y., Xu, C.S., Hotamışlıgil, G.S., et al. (2024). Spatial mapping of hepatic ER and mitochondria architecture reveals zonated remodeling in fasting and obesity. Nat. Commun. 15. 10.1038/s41467-024-48272-7.

21. Arruda, A.P., Pers, B.M., Parlakgül, G., Güney, E., Inouye, K., and Hotamisligil, G.S. (2014). Chronic enrichment of hepatic endoplasmic reticulum-mitochondria contact leads to mitochondrial dysfunction in obesity. Nat. Med. 20, 1427–1435. 10.1038/nm.3735.

22. Colleluori, G., Perugini, J., Di Vincenzo, A., Senzacqua, M., Giordano, A., and Cinti, S. (2022). Brown Fat Anatomy in Humans and Rodents. Methods Mol. Biol. 2448, 19–42. 10.1007/978-1-0716-2087-8_2.

23. Chen, Y., Zeng, X., Huang, X., Serag, S., Woolf, C.J., and Spiegelman, B.M. (2017). Crosstalk between KCNK3-Mediated Ion Current and Adrenergic Signaling Regulates Adipose Thermogenesis and Obesity. Cell 171, 836–848.e13. 10.1016/j.cell.2017.09.015.

24. Leaver, E. V., and Pappone, P.A. (2002). β-adrenergic potentiation of endoplasmic reticulum Ca2+ release in brown fat cells. Am. J. Physiol. Cell Physiol. 282. 10.1152/AJPCELL.00204.2001.

25. Parlakgül, G., Arruda, A.P., Pang, S., Cagampan, E., Min, N., Güney, E., Lee, G.Y., Inouye, K., Hess, H.F., Xu, C.S., et al. (2022). Regulation of liver subcellular architecture controls metabolic homeostasis. Nature 603, 736–742. 10.1038/s41586-022-04488-5.

26. Wikstrom, J.D., Mahdaviani, K., Liesa, M., Sereda, S.B., Si, Y., Las, G., Twig, G., Petrovic, N., Zingaretti, C., Graham, A., et al. (2014). Hormone-induced mitochondrial fission is utilized by brown adipocytes as an amplification pathway for energy expenditure. EMBO Journal 33, 418–436. 10.1002/embj.201385014.

27. Cannon, B., Jacobsson, A., Rehnmark, S., and Nedergaard, J. (1996). Signal transduction in brown adipose tissue recruitment: Noradrenaline and beyond. Int. J. Obes. 20.

28. Cero, C., Lea, H.J., Zhu, K.Y., Shamsi, F., Tseng, Y.H., and Cypess, A.M. (2021). β3-Adrenergic receptors regulate human brown/beige adipocyte lipolysis and thermogenesis. JCI Insight 6. 10.1172/jci.insight.139160.

29. Chang, C.L., Hsieh, T.S., Yang, T.T., Rothberg, K.G., Azizoglu, D.B., Volk, E., Liao, J.C., and Liou, J. (2013). Feedback regulation of receptor-induced ca2+ signaling mediated by e-syt1 and nir2 at endoplasmic reticulum-plasma membrane junctions. Cell Rep. 5, 813–825. 10.1016/j.celrep.2013.09.038.

30. Emrich, S.M., Yoast, R.E., and Trebak, M. (2022). Physiological Functions of CRAC Channels. Annu. Rev. Physiol. 84, 355–379. 10.1146/annurev-physiol-052521-013426.

31. Lewis, R.S. (2020). Store-operated calcium channels: From function to structure and back again. Cold Spring Harb. Perspect. Biol. 12. 10.1101/cshperspect.a035055.

32. Xue, K., Wu, D., Wang, Y., Zhao, Y., Shen, H., Yao, J., Huang, X., Li, X., Zhou, Z., Wang, Z., et al. (2022). The mitochondrial calcium uniporter engages UCP1 to form a thermoporter that promotes thermogenesis. Cell Metab. 34, 1325–1341.e6. 10.1016/j.cmet.2022.07.011.

33. Chang, C.L., Chen, Y.J., Quintanilla, C.G., Hsieh, T.S., and Liou, J. (2018). EB1 binding restricts STIM1 translocation to ER-PM junctions and regulates store-operated Ca2+ entry. Journal of Cell Biology 217, 2047–2058. 10.1083/jcb.201711151.

34. Arruda, A.P., Pers, B.M., Parlakgul, G., Güney, E., Goh, T., Cagampan, E., Lee, G.Y., Goncalves, R.L., and Hotamisligil, G.S. (2017). Defective STIM-mediated store operated Ca2+ entry in hepatocytes leads to metabolic dysfunction in obesity. Elife 6. 10.7554/eLife.29968.

35. 35. Orci, L., Ravazzola, M., Le Coadic, M., Shen, W.W., Demaurex, N., and Cosson, P. (2009). STIM1-induced precortical and cortical subdomains of the endoplasmic reticulum. Proc. Natl. Acad. Sci. U. S. A. 106, 19358–19362. 10.1073/PNAS.0911280106.

36. 36. Grigoriev, I., Gouveia, S.M., van der Vaart, B., Demmers, J., Smyth, J.T., Honnappa, S., Splinter, D., Steinmetz, M.O., Putney, J.W., Hoogenraad, C.C., et al. (2008). STIM1 Is a MT-Plus-End-Tracking Protein Involved in Remodeling of the ER. Current Biology 18, 177–182. 10.1016/J.CUB.2007.12.050.

37. Hall, D.D., Takeshima, H., and Song, L.S. (2024). Structure, Function, and Regulation of the Junctophilin Family. Annu. Rev. Physiol. 86, 123–147. 10.1146/annurev-physiol-042022-014926.

38. Behrens, J., Braren, I., Jaeckstein, M.Y., Lilie, L., Heine, M., Sass, F., Sommer, J., Silbert-Wagner, D., Fuh, M.M., Worthmann, A., et al. (2024). An efficient AAV vector system of Rec2 serotype for intravenous injection to study metabolism in brown adipocytes in vivo. Mol. Metab. 88. 10.1016/j.molmet.2024.101999.

39. Maus, M., Cuk, M., Patel, B., Lian, J., Ouimet, M., Kaufmann, U., Yang, J., Horvath, R., Hornig-Do, H.T., Chrzanowska-Lightowlers, Z.M., et al. (2017). Store-Operated Ca2+ Entry Controls Induction of Lipolysis and the Transcriptional Reprogramming to Lipid Metabolism. Cell Metab. 25, 698–712. 10.1016/j.cmet.2016.12.021.

40. Plate, L., Cooley, C.B., Chen, J.J., Paxman, R.J., Gallagher, C.M., Madoux, F., Genereux, J.C., Dobbs, W., Garza, D., Spicer, T.P., et al. (2016). Small molecule proteostasis regulators that reprogram the ER to reduce extracellular protein aggregation. Elife 5. 10.7554/eLife.15550.

41. Giacomello, M., Pyakurel, A., Glytsou, C., and Scorrano, L. (2020). The cell biology of mitochondrial membrane dynamics. Nat. Rev. Mol. Cell Biol. 21, 204–224. 10.1038/s41580-020-0210-7.

42. Leitner, B.P., Huang, S., Brychta, R.J., Duckworth, C.J., Baskin, A.S., McGehee, S., Tal, I., Dieckmann, W., Gupta, G., Kolodny, G.M., et al. (2017). Mapping of human brown adipose tissue in lean and obese young men. Proc. Natl. Acad. Sci. U. S. A. 114, 8649–8654. 10.1073/pnas.1705287114.

43. Shimizu, I., Aprahamian, T., Kikuchi, R., Shimizu, A., Papanicolaou, K.N., MacLauchlan, S., Maruyama, S., and Walsh, K. (2014). Vascular rarefaction mediates whitening of brown fat in obesity. Journal of Clinical Investigation 124, 2099–2112. 10.1172/JCI71643.

44. Tharp, K.M., Kang, M.S., Timblin, G.A., Dempersmier, J., Dempsey, G.E., Zushin, P.J.H., Benavides, J., Choi, C., Li, C.X., Jha, A.K., et al. (2018). Actomyosin-Mediated Tension Orchestrates Uncoupled Respiration in Adipose Tissues. Cell Metab. 27, 602–615.e4. 10.1016/j.cmet.2018.02.005.

45. Chen, Y., Zeng, X., Huang, X., Serag, S., Woolf, C.J., and Spiegelman, B.M. (2017). Crosstalk between KCNK3-Mediated Ion Current and Adrenergic Signaling Regulates Adipose Thermogenesis and Obesity. Cell 171, 836–848.e13. 10.1016/j.cell.2017.09.015.

46. Burkewitz, K., Feng, G., Dutta, S., Kelley, C.A., Steinbaugh, M., Cram, E.J., and Mair, W.B. (2020). Atf-6 Regulates Lifespan through ER-Mitochondrial Calcium Homeostasis. Cell Rep. 32. 10.1016/j.celrep.2020.108125.

47. Thuerauf, D.J., Hoover, H., Meller, J., Hernandez, J., Su, L., Andrews, C., Dillmann, W.H., McDonough, P.M., and Glembotski, C.C. (2001). Sarco/endoplasmic reticulum calcium ATPase-2 expression is regulated by ATF6 during the endoplasmic reticulum stress response: Intracellular signaling of calcium stress in a cardiac myocyte model system. Journal of Biological Chemistry 276, 48309–48317. 10.1074/JBC.M107146200.

48. Okuma, H., Saito, F., Mitsui, J., Hara, Y., Hatanaka, Y., Ikeda, M., Shimizu, T., Matsumura, K., Shimizu, J., Tsuji, S., et al. (2016). Tubular aggregate myopathy caused by a novel mutation in the cytoplasmic domain of STIM1. Neurol. Genet. 2, e50. 10.1212/NXG.0000000000000050.

49. Schiaffino, S. (2012). Tubular aggregates in skeletal muscle: Just a special type of protein aggregates? Neuromuscular Disorders 22, 199–207. 10.1016/j.nmd.2011.10.005.

50. Seale, P., Bjork, B., Yang, W., Kajimura, S., Chin, S., Kuang, S., Scimè, A., Devarakonda, S., Conroe, H.M., Erdjument-Bromage, H., et al. (2008). PRDM16 controls a brown fat/skeletal muscle switch. Nature 454, 961–967. 10.1038/NATURE07182.

51. Guney, E., Arruda, A.P., Parlakgul, G., Cagampan, E., Min, N., Lee, G.Y., Greene, L., Tsaousidou, E., Inouye, K., Han, M.S., et al. (2021). Aberrant Ca2+signaling by IP3Rs in adipocytes links inflammation to metabolic dysregulation in obesity. Sci. Signal. 14. 10.1126/scisignal.abf2059.

52. Roberts, E.J., Chavez, T., Hexemer, A., and Zwart, P.H. (2024). DLSIA: Deep Learning for Scientific Image Analysis. J. Appl. Crystallogr. 57, 392–402. 10.1107/S1600576724001390.

53. 53. Gallusser, B., Maltese, G., Di Caprio, G., Vadakkan, T.J., Sanyal, A., Somerville, E., Sahasrabudhe, M., O’connor, J., Weigert, M., and Kirchhausen, T. (2023). Deep neural network automated segmentation of cellular structures in volume electron microscopy. J. Cell Biol. 222. 10.1083/JCB.202208005.

54. Vincent, A.E., White, K., Davey, T., Philips, J., Ogden, R.T., Lawess, C., Warren, C., Hall, M.G., Ng, Y.S., Falkous, G., et al. (2019). Quantitative 3D Mapping of the Human Skeletal Muscle Mitochondrial Network. Cell Rep. 26, 996–1009.e4. 10.1016/j.celrep.2019.01.010.

55. Fischer, C.A., Besora-Casals, L., Rolland, S.G., Haeussler, S., Singh, K., Duchen, M., Conradt, B., and Marr, C. (2020). MitoSegNet: Easy-to-use Deep Learning Segmentation for Analyzing Mitochondrial Morphology. iScience 23. 10.1016/j.isci.2020.101601.

56. Lu, P.D., Jousse, C., Marciniak, S.J., Zhang, Y., Novoa, I., Scheuner, D., Kaufman, R.J., Ron, D., and Harding, H.P. (2004). Cytoprotection by pre-emptive conditional phosphorylation of translation initiation factor 2. EMBO J. 23, 169–179. 10.1038/SJ.EMBOJ.7600030.

57. Cannon, B., and Nedergaard, J. (2008). Studies of thermogenesis and mitochondrial function in adipose tissues. Methods in Molecular Biology 456, 109–121. 10.1007/978-1-59745-245-8_8.

